# Predicting the Frequency of Drug Side effects

**DOI:** 10.1101/594465

**Authors:** Diego Galeano, Alberto Paccanaro

**Affiliations:** Department of Computer Science, Centre for Systems and Synthetic Biology, Royal Holloway, University of London, Egham Hill, Egham, UK

## Abstract

Drug side effects are a leading cause of morbidity and mortality. Currently, the frequency of drug side effects is determined experimentally during human clinical trials through placebo-controlled studies. Here we present a novel framework to computationally predict the frequency of drug side effects. Our algorithm is based on learning a latent variable model for drugs and side effects by matrix decomposition. Extensive evaluations on held out test sets show that the frequency class is predicted with 67.8% to 94% accuracy in the neighborhood of the correct class. Evaluations on prospective data confirm the commonly held hypothesis that most post-marketing side effects are very rare in the population, with occurrences of less than 1 in a 10,000. Importantly, our model provides explanations of the biology underlying drug side effect relationships. We show that the drug latent representations in our model are related to distinct anatomical drug activities and that the similarity between these representations are predictive of the drug clinical activity as well as drug targets.

**One summary sentence:** novel explainable machine learning algorithm predicts the frequency of drug side effects in the population

## Introduction

The frequency of a drug side effect is one of the key variables in drug safety risk assessment (*1*), and it is crucial in the benefit-risk assessment for the clinical practice (*2*). It is well recognized that many drug side effects are not observed during clinical drug development, thus remaining a leading cause of morbidity and mortality in health care, resulting in an annual loss of billions of dollars (*3, 4*). This is due to the fact that even rigorous randomized, double-blind, placebo-controlled trials have limitations such as cohort size, time frame and lack of accrual (*5*). Thus, many rare or common delayed side effects are only detected after the drug has been marketed, in the so-called postmarking phase. Inaccurate estimation of drug side effect frequencies also constitutes a major risk for pharmaceutical companies as it can lead to drugs being withdrawn from the market (*6, 7*), or to a reassessment of side effect frequencies through new clinical studies, with high associated costs (*8*).

In this paper, we propose a novel method to predict the frequency of drug side effects. Given a few experimentally determined side effects, our method is able to predict the frequency of further side effects. This means that early clinical trials can be used to set the direction of the risk assessment before (or after) a drug enters the market. Our method can also be used in other aspects of clinical trial design, e.g. in the estimation of cohort size.

To our knowledge, this is the first computational method that successfully addresses the problem of predicting the frequency of drug side effects. Existing methods typically phrase the problem as a binary classification (*9-15*), where the aim is to predict the presence or absence of a drug side effect, not its frequency. One group of methods relies on the chemical structure of the compound or on the known drug target associations to produce de novo predictions (*11, 16, 17*). Pauwels et al. (*16*), for example, used sparse canonical correlation analysis to associate drug side effect profiles with their chemical substructure profiles and then predicted side effects for compounds with this model. Another group of methods exploits the structure of networks built by connecting drugs to known side effects (*9, 10, 15*). Cami et. al. (*9*), for example, built a bipartite drug side effect network and extracted feature covariates to learn a Bernoulli expectation model based on multivariate logistic regression.

Our idea to predict the frequency of side effects is to use a matrix decomposition model to learn a low dimensional latent representation of drugs and side effects that encodes the interplay between them. Our model is inspired by movie recommendation systems (*18-20*) that recommends movies to users: our recommendation system recommends side effects to drugs. Importantly, we constrain our matrix decomposition to be *non-negative*. This has the important advantage of making explicit the parts-based representation thus offering biological interpretability. In other words, we can interpret the latent variables in terms of the human anatomical system thus explaining the frequency predictions. Furthermore, drug latent representations can be used for predicting their clinical and molecular activity.

## Results

### Signature model of drug side effect frequencies

Five frequency classes are commonly used in clinical trials to describe the occurrence of drug side effects in clinical cohorts (Supplementary Note 1, Fig. S1). By coding these classes with integers between 1 and 5 (*very rare=1, rare=2, infrequent=3, frequent=4, very frequent=5*) we created an *(nxm)* matrix *R*, with 759 drugs (rows) and 994 unique side effect (columns) containing 37,441 frequency class associations (see Methods and Table S1). The remaining entries of the matrix were filled with zeros. The analysis of this dataset showed that drug side effects follow a long-tailed distribution (see Pareto distribution fit (Fig. S2) and QQ plot (Fig. S3)), where 80% of the associations are due to only about 30% of the side effect terms (Fig. 1a). Popular side effects, such as nausea or headache, account for most of the non-zero entries in the matrix and this implies that these popular side effects are reported on most drugs. Figure 1b shows that the distribution of frequency classes in our gold standard is *zero-inflated*: about 95% of the associations are unobserved. The average frequency rating value is 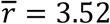, indicating a bias in clinical reports towards frequent side effects, that can be possibly attributed to the limitation of clinical studies at detecting less frequent side effects occurrences.

**Figure 1.**
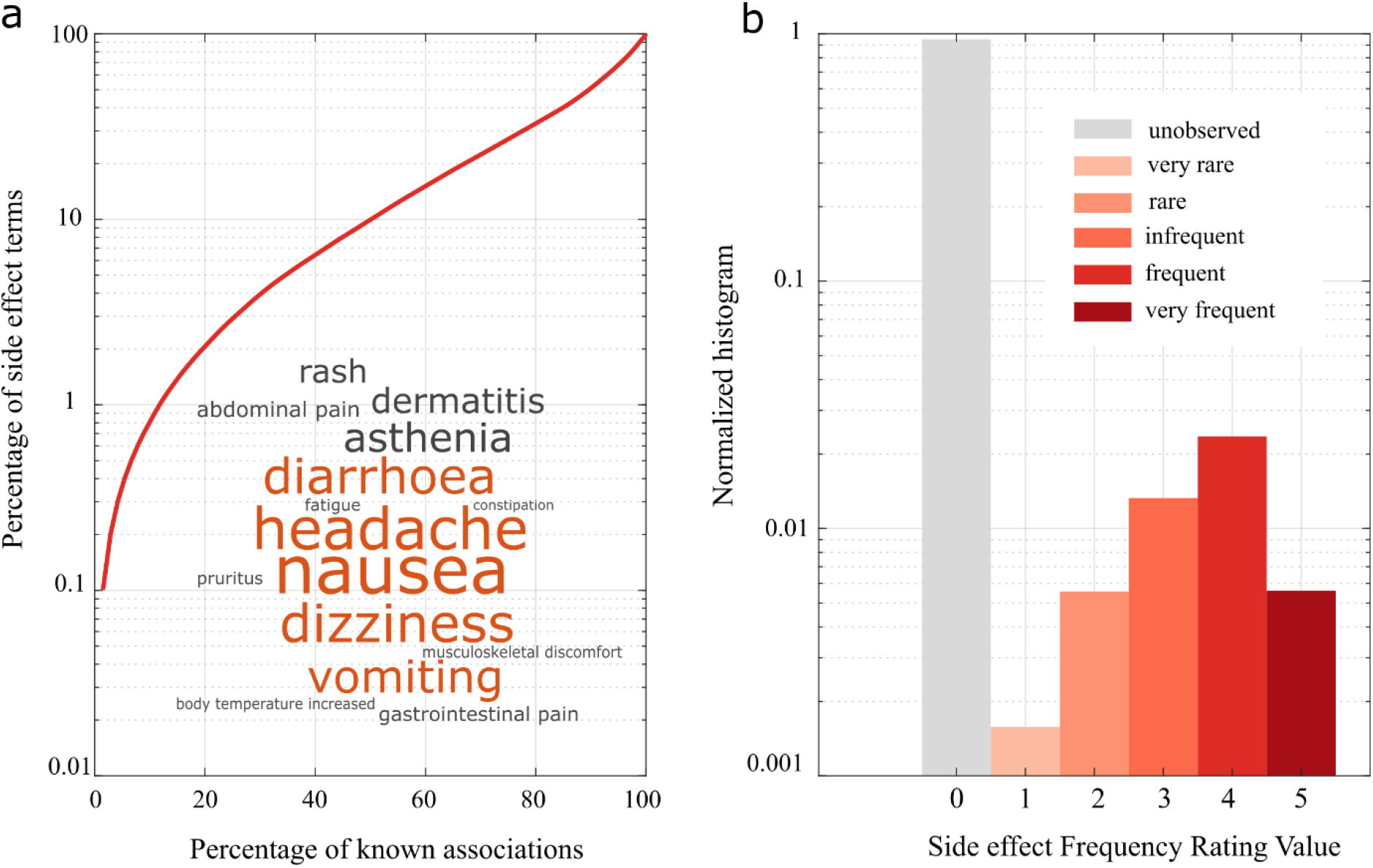
Distribution of drug side effects. The distributions are shown for the dataset **(a)** Side effects are ordered in decreasing order of popularity (number of drugs in which a side effect appear). *Inset*. A word cloud shows the fifteen most popular side effects. The size of the word is proportional to its popularity and the five most popular side effects amongst drugs are colored in orange **(b)** Histogram of drug side effect frequency values. The relative frequency of the population affected by drug side effects can be: very rare (less than 1 in 10,000), rare (1 in 10,000 to 1 in 1,000), infrequent (1 in 1,000 to 1 in 100), frequent (1 in 100 to 1 in 10) or very frequent (greater than 1 in 10) – shown in shaded red bars. The remaining of the associations are unobserved (grey bar).

The distribution resembles the one of the ratings found in movie datasets such as Netflix or MovieLens (*21*), where the 30% most popular movies account for 80% of the ratings; the distribution of rating values is bias towards high values (Fig. S4). An important group of methods for movie recommendation is based on matrix decomposition (*22*). The fundamental assumption is that both user and movies can be represented as vectors in a low-dimensional space (i.e. a small set of latent variables) and that a rating value for a specific *(user,movie)* pair can be obtained by the scalar product of the corresponding feature vectors. This assumption is reasonable for movie datasets where the latent variables can be thought of as modeling both movies genres and user preferences, such as thriller, horror, romance, etc.

This assumption is also reasonable for our problem: drugs and side effects can be represented as vectors in a low-dimensional space where the latent variables might model specific molecular mechanisms that elicit certain side effects (*23*). Therefore, our aim is to learn a low-dimensional representation for each drug (that we shall call *drug signature*, 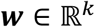), and a low-dimensional representation for each side effect (*side effect signature*, 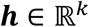), such that the frequency of a drug-side effect pair is modelled by the scalar product of the two feature vectors (illustrative example in Fig. S5). It amounts to decomposing our original matrix *R* into a product of two matrices, *W* and *H*, where the rows of *W* and columns of *H* are the signatures of drugs and side effects respectively; that is, *R_n×m_* ≈ *W_n×k_H_k×m_*, where *k* ≪ min(*n, m*) (see Methods). Our algorithm learns the matrices *W* and *H* by minimizing the following loss function:

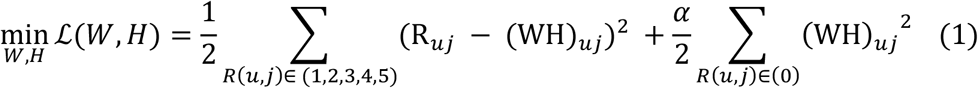

Subject to the non-negative constraints *W, H* ≥ 0

Here, the first summation aims at reconstructing *R* for the known class values, while the second one aims at reconstructing the zeros found in *R*. These two types of values need to be treated separately because they have inherently different meanings. While rating values {1, 2, 3, 4, 5} correspond to experimentally measured frequency of side effects, a zero entry simply indicates that no side effect was detected for a certain drug – which could either mean that the drug does not cause the side effect, or that it does, but it could not be detected^1^. The parameter *α*, ranging between 0 and 1, is set through cross-validation (see Methods), and controls the relative importance of the zeros; in other words, it represents our confidence in their correctness (Fig. S6). It should also be noted that the second summation also acts as a regularization factor, so no additional regularization terms are needed (see Supplementary Note 3). Finally, we impose non-negative constraints on our solution as it leads to a parts-based representation (*24*) and consequently to an increased interpretability of drug and side effect signatures, since only additive combinations of latent representations are allowed.

To minimize this function subject to the non-negative constraints, we developed an efficient iterative algorithm that uses a simple multiplicative update rule. Our procedure does not require setting a learning rate nor applying a projection function and satisfies the Karush-Kuhn-Tucker (KKT) complementary conditions of convergence (see Methods and Supplementary Note 5).

### Evaluation of the performance at predicting side effect frequency

We held-out 10% of the observed values in *R* for testing. The remaining 90% were used in a tenfold cross-validation procedure to set the two parameters of our algorithm: *k* (the number of latent features) and *α* (confidence in the zeros). During cross validation, performance was assessed using the root mean squared error (RMSE) with respect to the known associations for the five frequency classes, as well as the area under the receiver operating curve (AUROC) obtained when predicting the presence/absence of the associations (see Methods). A good performance on the cross-validation sets was obtained with α = 0.05 and *k* = 10 (mean RMSE = 1.372 ± 0.021 and mean AUROC= 0.920 ± 0.003) – see Fig. S7. Note that our algorithm is quite robust with respect to the choice of the parameters α and *k* – the sensitivity analysis is shown in Fig. S8-S9.

On the held-out test set our model scored a RMSE of 1.32 and an AUROC of 0.923. Figure 2 (top) shows, for each of the five frequency classes in the test set, the histogram of the values that were predicted for that class. The Pearson correlation between the predicted scores and their corresponding frequency classes was *ρ* = 0.60 (Significance, *p* < 2.22*e* − 308); the differences between the distributions of scores for the five frequency classes were statistically significant. As a further verification of the effectiveness of our scores at capturing the frequency classes, we tested whether the same properties hold for scores calculated by the Predictive Pharmaco-safety Networks (PPN-NET) (*9*), which learns probabilities of drug side effect associations. We found that PPN-NET scores are only weakly correlated to the frequency of drug side effects (Pearson, *ρ* = 0.09, *p* < 3.75*e* − 08).

**Figure 2.**
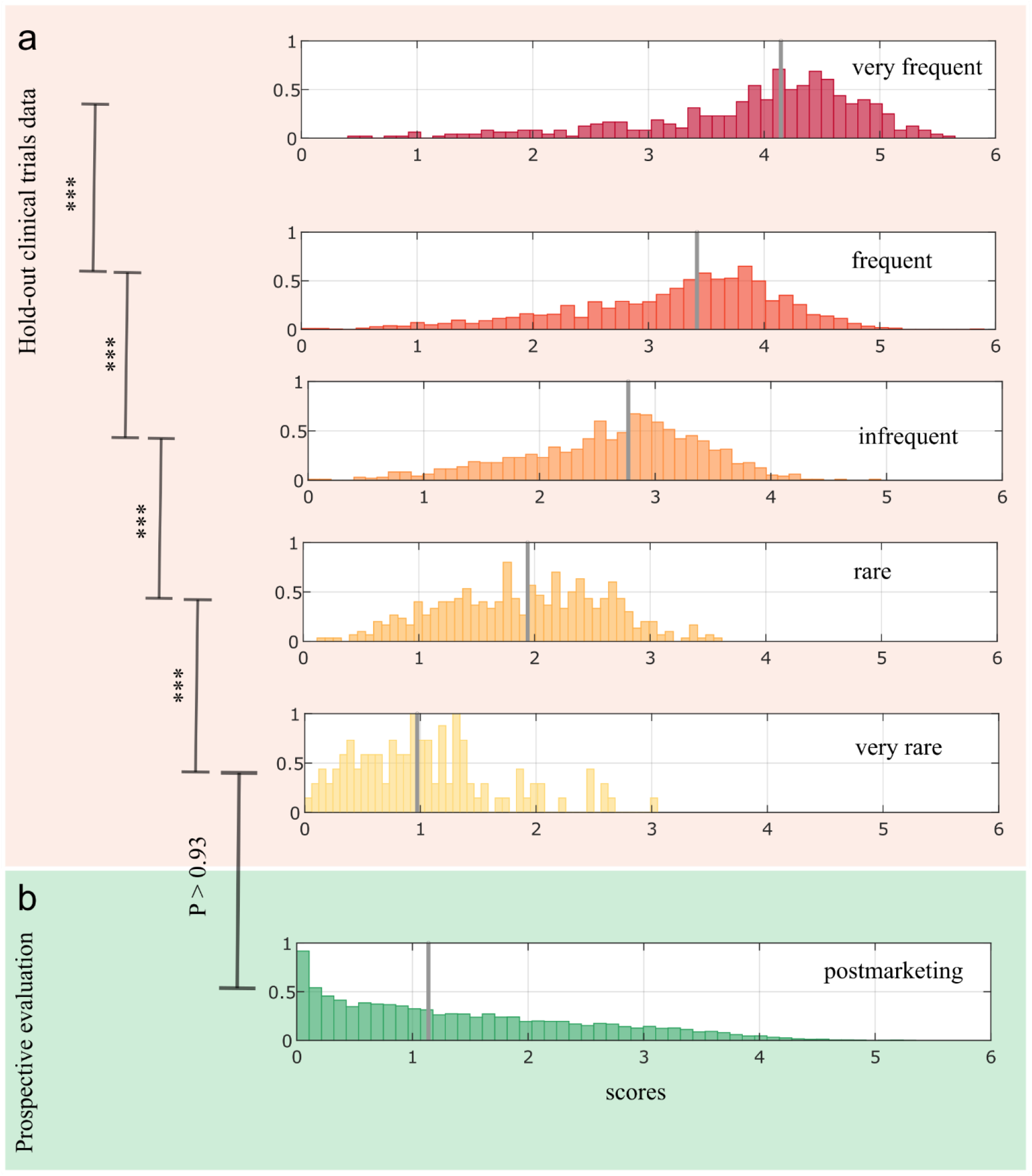
Distributions of scores for held out and post-marketing datasets. **(a)** Normalized histogram of scores obtained for each of the five frequency classes in the held out test set. The differences in the distributions between the classes are statistically significant: rare vs very rare (p < 2.80×10^−12^), infrequent vs rare (p < 1.31×10^−40^), frequent vs infrequent (p < 3.45×10^−51^) and very frequent vs frequent (p < 9.00×10^−26^). The smoothness of the histogram is related to the class numerosity. **(b)** Normalized histogram of scores obtained for post-marketing reports. There are no significant differences between these scores and those obtained for very rare side effects. Significance levels between the scores are indicated with asterisks (p ≤ 0.001, ***), (p ≤ 0.01, **). Two-tailed Wilcoxon rank sum test was used all the cases. Median values are shown as grey vertical lines.

To predict the frequency class of a given drug-side effect pair, we need a way to assign scores to frequency classes. To do this, we learned the likelihood function of each class (i.e.*P*(*x*|*class*)) and assigned scores to classes by maximum likelihood. Here, the likelihood functions were learned on the cross-validation sets (see Methods). Note that, due to biases in the dataset, we cannot obtain reasonable estimates for the priors (thus forcing us to use uniform priors for each class). Furthermore, due to the lack of experimentally validated zero values, in order to discriminate the zeros we followed similar approach of Cami *et al*. (*9*) and chose a threshold using the ROC curve at a sensitivity of 0.97 given a specificity of 0.57 (see Methods).

Figure 3a shows the percentage of accuracy at predicting frequency classes. For drug-side effect pairs in any given class, the most predicted class is the correct one. For each class, the prediction accuracy ranges from 55.2% to 75.5% when including the immediately lower class, and 67.8% to 94% when both adjacent classes are considered. Looking at the first column in the figure, we notice how the system very rarely (0.72%) fails to detect a side effect that is very frequent, and seldom misses side effects in the rare frequent (2.68%), infrequent (2.52%) and rare (3.11 %) classes. The number of undetected side effects only increases for the very rare class (16.94%) – this is probably due to the small number of known association in this class (see Figure 1b and Fig. S8), which in turn reflects the bias in clinical trial reports.

**Figure 3.**
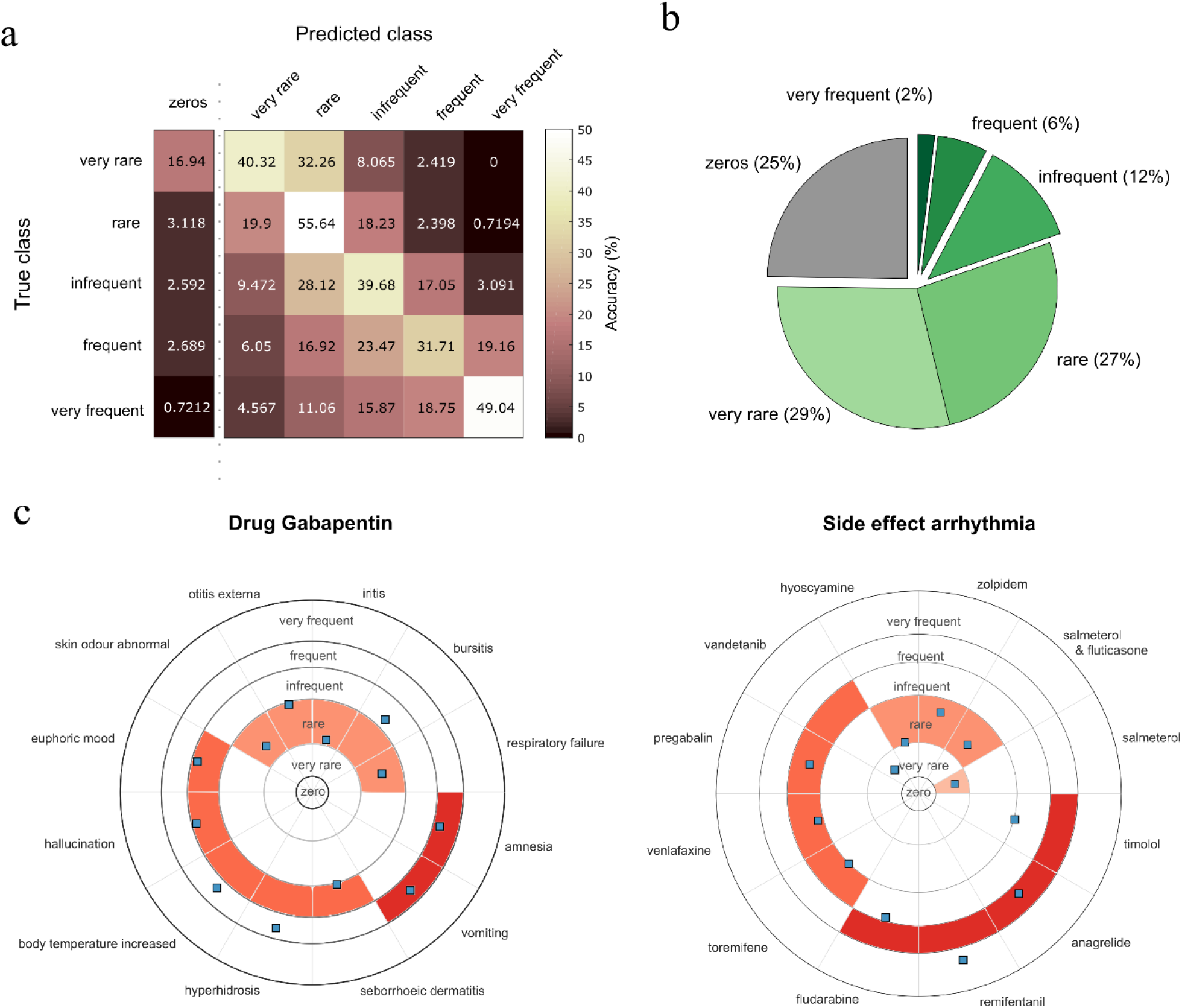
Evaluation of side effect frequency predictions. **(a)** Accuracy percentages for the predictions in the held-out test set. Frequency classes are predicted by maximum likelihood. Zeros, corresponding to “no side effect” prediction, are predicted for score values below 0.42 (corresponding to 0.97 sensitivity given 0.57 specificity). **(b)** Distribution of classes assigned to post-marketing data. **(c)** Illustrative example from the held-out test set. Twelve randomly chosen predictions for the anticonvulsant drug Gabapentin (*left*) and the cardiovascular side effect arrhythmia (*right*) are shown around polar plots, each in a dedicated sector. Gray concentric circles between frequency classes correspond to thresholds learned by maximum likelihood. The correct class for each association is colored in each circular sector while predicted scores are shown as blue squares.

As illustrative examples, Figure 3c (and Fig. S11-12) presents the predicted frequency scores for the anticonvulsant drug Gabapentin, a top fifty prescribed drug in the U.S. (*25*), and the cardiovascular side effect arrhythmia, important in cardiotoxicity assessment (*26*).

We further tested the performance of our system at predicting the frequency of side effects that were detected after the drugs had reached the market. Only the presence/absence of these so-called post-marketing associations are known and these are normally regarded as side effects of very rare occurrence in the population (*27, 28*). When we analyzed the distribution of scores obtained by our system for these associations, we found no significant differences between them and the scores of very rare associations in the held-out test set (Fig. 2 bottom and Fig.S7, Wilcoxon Significance, p > 0.936). Fig. 3b shows the percentages of post-marketing associations assigned to each class by maximum likelihood. It can be seen that 55.5% of these associations were predicted to be either very rare or rare, while only 2% were predicted as very frequent. Amongst these, the most recurrent side effects were gastrointestinal disorder (16/180), vomiting (7/180), nervous system disorders (7/180) and infection (7/180).

### Drug signature similarity predicts clinical response and drug targets

The effectiveness of our model at predicting the frequency of side effects prompted us to analyze whether the learned signatures are informative of the biology underlying drug activity.

We investigated the link between drug signatures and clinical responses. We hypothesized that the signature for two drugs should be more similar when they share clinical activity. Drugs were grouped based on their main Anatomical, Therapeutic and Chemical (ATC) class (Table S2), a hierarchical organization of terms describing clinical activity where lower levels of the hierarchy contain descriptors that are more specific. Figure 4a shows that, in most cases, the cosine similarity between the signature of drugs within an anatomical class is higher that the similarity between classes (Fig. S13, Supplementary Tables 5-6). We further proved that drug signature similarity is related to drug clinical activity similarity by showing that drug signatures similarity is predictive of shared ATC category at each of the different levels (Fig. 5b and Fig. S14). It is important to note that the prediction performance of the drug signature similarity increases as we consider terms located lower in the ATC hierarchy – this correctly reflects the fact that drug clinical responses become more similar as we move to lower (more specific) levels of the ATC hierarchy.

**Figure 4.**
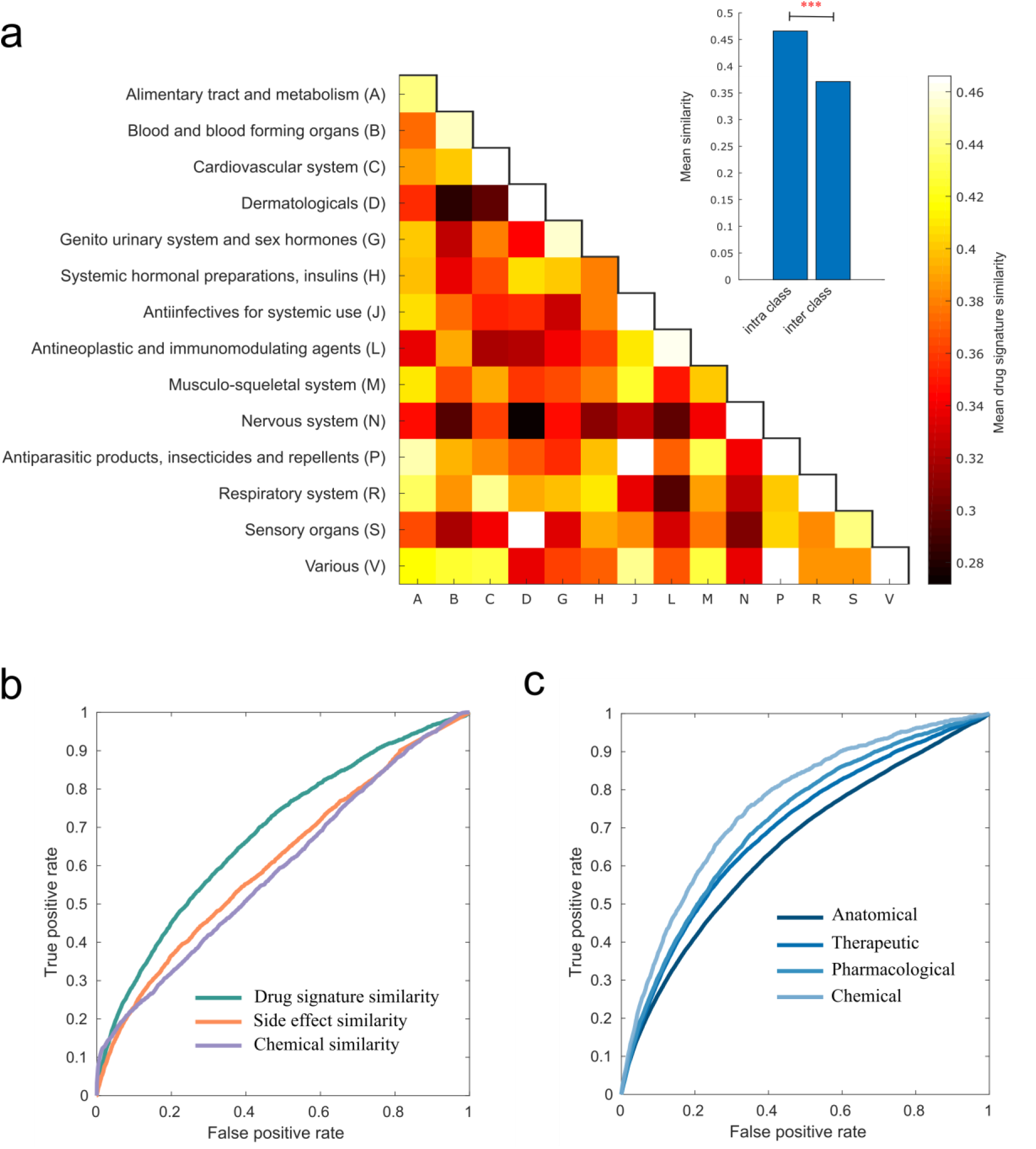
Drug signatures capture the drug clinical and molecular activity. **(a)** Heat maps of mean drug signature similarities per anatomical class. Each (x, y) tile represents, for each main Anatomical, Therapeutic and Chemical (ATC) drug category, the mean similarity of drug pairs where one drug belong to category x and the other to category y. The value ranges from 0.27 (Nervous system - Dermatological) to 0.55 (Nervous system-Nervous system). The colours range between the minimum mean similarity and 0.466, with all values above 0.466 (In the diagonal: 0.471 (C), 0.512 (D), 0.55 (N), 0.47 (P), 0.52 (R), 0.475 (V)) set to 0.466. *Inset*: the average intra-class similarity is significantly higher than the average interclass similarity (t-test p-value < 2.62e-13). **(b)** ROC curve representing the ability of the drug signature similarity, side effect similarity and Tanimoto chemical similarity scores to predict which pairs of drugs share targets. For 435 drugs in our dataset, 2,808 pairs were known to share molecular targets whereas 91,587 pairs were unknown. **(b)** ROC curve representing the ability of the drug signature similarity to predict which pairs of drugs share Anatomical, Therapeutic and Chemical (ATC) category at each of the different levels in the ATC taxonomy. Drug signature similarity was predictive of clinical drug activity at different levels: *anatomical class* (38,711 pairs share vs 248,950, AUROC = 65.33%), *therapeutic subclass* (11,960 pairs share vs 275,701, AUROC = 69.51%), *pharmacological subclass* (5,522 pairs share vs 282,139, AUROC = 71.54%) and *chemical subclass* (1,736 pairs share vs 285,925 do not, AUROC = 76.05%).

Encouraged by these results, we decided to test whether drug signature similarity can even be used for the prediction of drug targets. Having framed this as a binary classification problem (see Methods), we found that drug signature similarities are predictive of shared drug protein targets (AUROC = 68.38%) (Fig. 5a,) and performs better than baselines previously used elsewhere (*27, 29*), such as the 2D Tanimoto chemical similarity (AUROC = 59.26%) and to the Jaccard side effect similarity (AUROC = 61.07%). Note that the difference in the distribution of similarities between the two sets of drug pairs (those that share and those that do not share targets) is statistically significant (Wilcoxon Significance, p < 2.85×10^−242^) (see Fig. S14).

We also analyzed the link between side effect signatures and the anatomy and physiology of the side effect phenotypes (Fig. 4b and Fig. S15-16). Side effects were grouped based on their anatomical class according to MedDRA (Table S3). We found that signature for two side effects tend to be more similar when they are phenotypically related. Figure 4b shows that, in most cases, the cosine similarity between the signature of side effects within system organ classes (top level of the MedDRA hierarchy) is higher that the similarity between classes. Moreover, side effect signature is predictive of shared MedDRA class at each of the different levels and predictions improve as we move to more specific terms in the MedDRA hierarchy (see Fig. S17).

## Discussion

We presented a novel framework for predicting the frequency of drug side effects. Our model learns a low dimensional representation of drugs and side effects that encodes biologically meaningful information. We can envision the use of our system by safety professionals in pre-and post-marketing drug development. In premarketing, to assist controlled clinical trials design as a hypothesis generator based on previously collected associations from earlier phases of human trials. In post-marketing studies, to provide an alternative tool for surveillance reporting systems in the early discovery of rare side effects. Furthermore, our method might be useful to public health regulators such as the US Food and Drug Administration (FDA) in the assessment of drug safety.

Our method is inspired by collaborative filtering models from movie recommendation systems. However, our method differs from the standard movie recommendation system in the model assumption. While in movie recommendation unobserved values represent missing values, in our problem, they can also represent a zero value that indicates that the side effect was not detected for a certain drug. To account for this uncertainty, we have developed an objective function and a novel multiplicative algorithm with theoretical guarantees of convergence (see Supplementary Note 3). Other decomposition methods, such as singular value decomposition (SVD), or non-negative matrix factorization (NMF) (*30*), do not account for this uncertainty, and as such, they cannot weight for their importance in the learning.

Drug side effects have been previously found predictive of molecular targets (*23*) and therapeutic indications (*31*). One important question was whether our model’s signatures were associated with the molecular and the clinical drug activity. We found that drugs with similar signatures were more likely to share a protein target and to belong to the same anatomical, therapeutic, pharmacological and chemical class.

Biases in the dataset represents a challenge. We observed that the frequency of side effects from clinical trials are biased towards frequent side effects (Fig. 1b). Recent reports also indicate that clinical trials are biased towards Caucasians: 86% of clinical trials cohorts were white-dominated in 2014 (*32*). Numerous previous research also reported divergent drug responses in subjects with a different genetic background (*33*). It is possible to improve our framework applicability by integrating metadata from clinical trials. Additional knowledge about drugs or side effects can be incorporated into our model as constraints. For example, by considering gender or age-related side effects we might be able to enforce true zeros in the associations.

Our framework applicability requires known experimental associations for each drug and for each side effect from placebo-controlled studies. It would be interesting to extend the analysis presented here to predict the frequency of side effects for novel compounds based solely on chemical or cellular features. For instance, by exploiting similarities in chemical structure or in activity across cell lines. It can pave the way to establish direct relationships between chemical or cellular features and the phenotypic drug signatures of our model. Furthermore, drug signatures could be useful in the study of adverse effects produced by drug combinations, for instance, by studying whether additive drug signatures can explain the exponential risk increase of adverse effects.

## Material and Methods

### 1. Datasets

We used the drug side effect frequencies from the Side effect Resource (SIDER) database version 4.1 (*34, 35*). In the database, around 40% of the pairs have frequency information whereas, for the remaining associations, the frequency is unknown. Drugs are indexed by their PubChem IDs and side effect terms are mapped to the Medical Dictionary for Regulatory Activities (MedDRA) taxonomy. We only considered side effect terms that were Preferred-Terms (PT) in MedDRA. We also kept only the drugs with known monotherapy Anatomical Therapeutic and Chemical (ATC) classification according to the 2018 World Health Organization (WHO) release. Frequency data is presented in different formats, i.e. as exact, range or frequency label. We used frequency labels common in clinical trials, i.e. very rare, rare, infrequent, frequent and very frequent. Finally, we encoded the frequency labels using natural numbers from one to five (five-star rating system), i.e. very rare (=1), rare (=2), infrequent (=3), frequent (=4) and very frequent (=5). Our gold-standard dataset contains 37,441 frequency associations for 759 drugs and 994 side effect terms (see Supplementary Note 1).

We used Drugbank v5.0.5 (*36*) to extract drug protein targets and the SMILES fingerprints of drugs. The drug Anatomical,Therapeutic and Chemical (ATC) classes as well as the route of administrations (Adm.R) were obtained from WHO release 2018 (see Supplementary Note 2).

### 2. Matrix decomposition model

We modeled the drug side effect frequency estimation as linear combinations of drugs and side effects activation patterns over a set of latent features. To predict a drug-side effect frequency rating value, each component in the drug signature is multiplied by the corresponding component of the side effect signature and then the products are summed together. Thus the frequency of a side effect *j* for a given drug *d* can be expressed it as a combination of *k* components as follow,

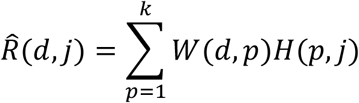

where *W*(*d,p*) is the drug signature and *H*(*p,j*) is the side effect signature on the *p^th^* component.

### 3. Objective function and the decomposition algorithm

Let *R_n×m_* be a matrix of n drugs and m side effects which contains the observed frequency ratings from clinical trials for each drug. For any matrix *A*, let the projection *A_R_* (both with dimension n × m) be such that:

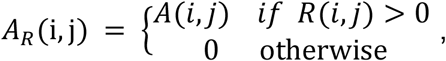

this amounts to obtaining all entries in matrix *A* that correspond to the non-zero entries in the matrix *R* (note that trivially *R_R_ = R*). Likewise, the negative projection of matrix *A* denoted by *A_¬R_* is defined as:

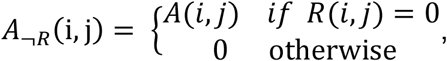

which amounts to obtaining all entries in matrix *A* that correspond to the **zero** entries in matrix *R*.

Using this notation, we can now write our objective function as:

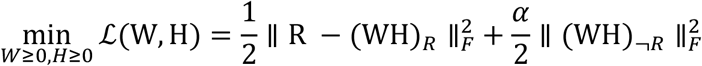

where *W_n×k_* contains the drug signatures, *H_k×m_* the side effect signatures, *α* is the confidence on the null associations, and 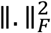 is the Frobenius norm. Notice that this objective is the matrix form of equation (*1*) in the main manuscript.

The objective function is composed as the sum of two terms:

- The term 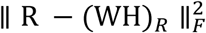 learns to fit the linear model 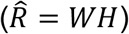 on the observed clinical trials data;
- The term 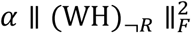 learns to fit the model to unobserved associations weighted by a confidence coefficient *α*.

We obtained a multiplicative update rule that minimizes 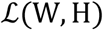 at each iteration without requiring a step size parameter nor a projection function, i.e.,

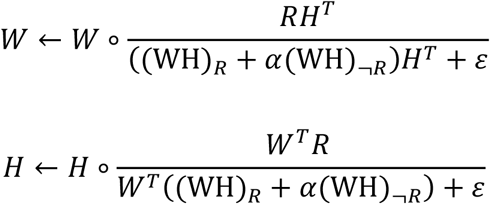

where ◦ denotes element-wise product, the division is element-wise, and *ε* = 10^−16^ is a small number added to prevent division by zero (*37*). In our experiments, both matrices are initialised as random dense matrices uniformly distributed in the range [0, σ], with σ = 0.1.

Furthermore, to avoid the well-known degeneracy (*24*) associated with the invariance *WH* under the transformation W → WΛ and H → Λ^−1^H, for a diagonal matrix Λ, we normalized *H* at each iteration as follow;

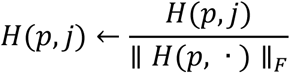

where *H*(*p*, ·) denotes the *p*-th row in *H*.

The convergence criteria for our algorithm was based on the termination tolerance on relative change in the elements of *W* and *H*. Default value was TolX < 10^−3^, that occurred in typically in about 2,000 iterations for *k* = 10.

The algorithm satisfies the Karush-Khun-Tucker (KKT) supplementary condition of local minima convergence (Proof in Supplementary Note 5). The code for the implementation of the algorithm is provided (Supplementary Code 1).

### 4. Cross-validation procedure and model selection

We used a tenfold cross-validation procedure (*22*) for the setting of the parameters *k* and *α*. Let Ω denote the set of observed entries in *R*. We first randomly removed 10% of the entries in Ω to create a hold-out test set E. The entries in T = Ω − E were then randomly split into ten disjoint subsets *s_j_*, j ∈ (1,…,10) for cross-validation. Then, for each fold *s_j_,* , the entries in *s_j_* were used for testing, while the remaining T − *s_j_* sets were used for training. The evaluation was computed as follow; (A) To evaluate the ability of the model for the accurate estimation of the frequency rating value, we use the Root Mean Squared Error (RMSE) on the unseen test set *s_j_*:

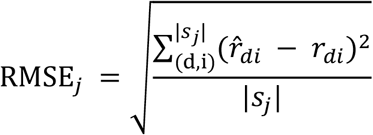

where |*s_j_*| denotes the cardinality of the test set, *r_di_* ∈ *s_j_* is the frequency rating value of drug side effect pair and 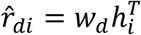 is the prediction for the dth drug and ith side effect.

(B) To evaluate the ability of the model to rank higher the known associations in *s_j_* with respect to the set of remaining unobserved entries M = ((d, j) ∉ T − *s_j_*), we used the Area Under the Receiver Operating Characteristic Curve (AUROC), a standard measure in binary classification and binary side effect prediction (*9*). We repeated this procedure for each of the sets in *s_j_* and then computed the mean RMSE and the mean AUROC for the ten folds. To select the model parameters, we first chose *α* based on a close to optimal binary classification performance (AUROC) while ensuring a good RMSE. We found that a good choice of *α* was 0.05. We then chose the optimal number of latent features that minimized the mean RMSE. This occurred for *k* = 10.

Finally, to show that the performance on a hold-out test set was as good as the ones from cross-validation, we trained our model 1,000 times (to find a good local minimum) with the chosen parameters using all the entries in Ω − E and tested it on the hold-out test set E.

### 5. Maximum likelihood estimate for frequency class prediction

To predict specific frequency classes, we first collected the predicted scores 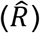 in the cross-validation for each of the values in each fold *s_j_*,j ∈ {1,…,10}. Then, for each of the five frequency classes, we fitted a normal kernel smoothing function to the predicted scores and obtained a probability density function (pdf) for each of the five classes. Finally, to evaluate the performance of the method on the hold-out test set E, we assigned each entry in E to the highest *P*(class|*x*) given a score *x* predicted for a drug side effect pair in E. To assign a hard threshold for the class zero (frequency zero), we used the ROC curve obtained in the test set E. We obtained a sensitivity of 0.97 given a specificity of 0.57. The threshold obtained for the class zero was 0.426. For the following classes, the obtained thresholds were 1.26, 2.43, 3.25 and 3.93.

### 6. Similarity between drug and side effect signatures

We quantify the similarity between two drug or side effect signatures using the cosine similarity over the set of latent representations. In detail, given two drug signatures *w*_1_ and *w*_2_ (each a vector of *k* dimensions), the drug signature similarity is given by the dot product of the vectors divided by the product of the norm of each vector.

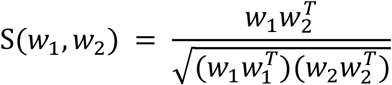

Therefore, the similarity for non-negative signatures ranges from 0 to 1.

### 7. Using drug signature similarity to predict clinical activity and drug targets

We used the drug signature similarities to predict whether two drugs share or not clinical activity or drug protein targets. Following the approach by *Tattoneti et al*. (*27*), we frame it as a binary classification problem, where the scores are the signature similarities and the labels are one if two drugs share activity and zero otherwise. The performance is measured using the AUROC. We applied the same procedure for side effect signatures.

## Acknowledgments

**Funding**: the work is supported in part by Biotechnology and Biological Sciences Research Council (BBSRC), grants BB/K004131/1, BB/F00964X/1 and BB/M025047/1 to A.P., CONACYT Paraguay Grants 14-INV-088 and PINV15-315, and NSF Advances in Bio Informatics grant 1660648.

## Author contributions

D.G. and A.P. conceived the study. D.G. devised and implemented the algorithm and conducted experiments. A.P. supervised the project. D.G. and A.P. discussed the results and implications. D.G. and A.P. wrote the manuscript.

## Footnotes

1 This is different from what happens in movie recommender systems where zeros correspond to missing values only – see Supplementary Note 4 for details.

