## Supplementary Materials for "Predicting the Frequency of Drug Side effects"

**This PDF file includes:**

Supplementary Notes 1-5

Supplementary Figures 1-26

Supplementary Tables 1-3

### Supplementary Notes

#### Supplementary Note 1: Gold standard drug-side effect frequency data

We started with 1,556 marketed drugs listed in the Side effect Resource Database (SIDER) 4.1 (*1*) that contains 4,251 side effect terms mapped in the Medical Dictionary for Regulatory Activities (MedDRA) v20.0. In SIDER 4.1, around 40% of the drug-side effect pairs contains frequency information, whereas for the remaining 60% the frequency is unknown. Side effect terms were annotated with their MedDRA lowest level term (LLT) and their preferred term (PT). One or more LLTs are equivalent to the same PT. For instance, the LLTs Creatinine increased (C0151578), Blood creatinine increased (C0235431), Serum creatinine increased (C0700225) and Plasma creatinine increased (C0858118) corresponds to the same MedDRA PT Blood creatinine increased (C0235431). Thus, we used only side effect MedDRA PT associations to avoid redundancies. For each drug, we retrieved the associated side effect frequencies. Side effect frequencies were listed as exact frequencies, e.g. 1%, range of frequencies, e.g. 2-5% or as frequency class e.g. very rare, rare, infrequent, frequent and very frequent. Placebo frequency was also available either as range or exact frequency. In Fig. S1 we summarize the types frequency formats in a Venn diagram depicting the three different sets of data formats. For a given drug- side effect pair, multiple frequencies might also be available, for example, from clinical trials for different indications (*2*). For convenience, we standardize all the frequencies into frequency classes. Thus, we standardized exact frequencies to frequency classes using the equivalent range of occurrences reported in clinical trials (Table S1) which is regulated by the Council for International Organizations of Medical Sciences (CIOMS). For the different subsets depicted in Figure S1, we pre-processed the frequency data as follow;

1. **Subset A - (A ∪ B)**. When only the exact or range frequency was available we compute the median frequency and then map to a frequency class.
2. **Subset B - (A ∪ C)**. When only the frequency class label was available we kept the labels but normalized the following terms: very rare, rare, infrequent (or uncommon), frequent (or common) and very frequent (or very common).
3. **Subset C - (A ∪ B)**. We have found 215 pairs for which the placebo frequency was found but the drug frequency was not listed in SIDER 4.1. We checked manually several pairs to confirm that the drug frequency was indeed missing in the database. We discarded these pairs.
4. **Subset A ∩ C**. We only retained pairs for which the median frequency in the intervention cohort was higher than the median in the placebo cohort. For 2,474 pairs, the median frequency in both groups were either comparable or the placebo frequency was higher than the drug frequency (403 associations). Previous studies indicates that these associations are likely to be cause by the disease or by the so-called nocebo effect (*3*), i.e. patients that anticipate a side effect on a medication are more likely to reported it (*1*). We discarded these pairs to avoid possible confounders in the associations.
5. **Subset A ∩ B**. We compute the median frequency value and map to a frequency class. We also kept the frequency class from B.
6. **Subset B ∩ C**. We discarded the placebo frequencies for which not intervention frequency was found. We kept the frequency class from B.

7. **Subset A ∩ B ∩ C**. We retained pairs for which the median frequency was higher in the intervention cohort than the placebo cohort. We also kept the frequency class from B.

We then encoded the side effect frequency classes using a five-star rating system (Table S1), where each frequency class is assigned to a natural number from one to five, representing the less to the most frequent side effects. We have also found that for around 13% of the pairs, we could have more than one rating value for a given drug side effect due to the multiple intersections in the data. 8% of these frequency classes were inconsistent. These might be due to clinical trials from different indications for the same drug (*1*). For these cases, we average the rating values and then round them to the nearest highest integer. Until here, we have extracted 41,546 frequency associations for 860 drugs and 1,011 side effects. Furthermore, we kept only drugs with known monotherapy Anatomical Therapeutic and Chemical (ATC) category according to the 2018 World Health Organization (WHO) release and side effects with known MedDRA category of disorders (Supplementary Table S3). In total, our gold-standard dataset contains 759 drugs with 994 side effect terms with 37,441 known rating values.

#### Supplementary Note 2: Additional datasets

##### Binary drug side effect associations

For the gold-standard dataset of 759 marketed drugs with 994 side-effect terms, we retrieved the binary drug-side effect associations with unknown frequencies from SIDER 4.1. We found 55,382 binary associations in SIDER. These binary associations are either from clinical trials with unreported frequency or from post-marketing reports added to drug leaflets (*1*). In fact, 9,387 pairs (16.94 %) were found with an associated post-marketing label. We used this subset of binary drug side effect associations as prospective evaluation set and to analyse the predicted frequencies. We refer to this as *post-marketing* set.

##### Drug molecular targets

We retrieved the known drug-target interactions from DrugBank release 5.0.5 (2016-08-17) (*4*). We mapped the drugs from SIDER to DrugBank using the PubChem IDs and the mapping provided in DrugBank. We retrieved molecular targets (with known or unknown pharmacological action) for 435 drugs from our gold-standard dataset. In total, 590 unique protein targets were found for the set of drugs with 1,759 known associations.

##### Drug chemical fingerprints

We retrieved the known drug SMILES fingerprint from DrugBank release 5.0.5 (2016-08-17) (*4*). We found 442 drugs from our gold-standard with SMILES. We then computed the 2D Tanimoto chemical similarity based on the fingerprint using the Open-Source Cheminformatics (RDKit) (*5*) in python.

**Drug Route of Administration (Adm.R)**

We retrieved the route of administration of each drug for each associated monotherapy ATC category from WHO 2018 release (Supplementary Database S2). The routes of administration of the drugs can be implant, inhalation, instillation, nasal (N), oral (O), parenteral (P), rectal (R), sublingual/buccal/oromucosal (SL), transdermal (TD) and vaginal (V).

#### Supplementary Note 3: On the data-driven regularization and possible variations of the method

*The notation here follows the notation presented in the Methods section of the main paper.*

The two most important assumptions in our model are that *(i)* the frequency of drug side effect pair can be modelled as the scalar product of their low-dimensional feature vectors and (ii) the confidence on the unobserved drug side effect associations (zeros) should be set by cross-validation. To accomplish this, we learned from the observed (5% associations) and unobserved (95% associations) entries with a separate confidence. The first term in our objective function learns the representations that fits the observed associations, whereas the second term in our objective function $\alpha \parallel (\mathrm{WH})_{\neg R} \parallel_F^2$ can be interpreted as a data-driven regularization. To illustrate this point, we can write our objective function as follow,

$$\min_{W \geq 0, H \geq 0} \mathcal{L}(\mathrm{W}, \mathrm{H}) = \frac{1}{2} \parallel \mathrm{R} - (\mathrm{WH})_R \parallel_F^2$$

$$\text{Subject to} \quad \sum_{(d,j) \notin \Omega} \hat{\mathrm{r}}_{dj}^{\ 2} \leq \delta$$

where $\delta \geq 0$ is a regularization parameter closely related to $\alpha$, its corresponding Lagrange form. Here Ω denotes the set of observed entries in R. The constraint $\sum_{(d,j) \notin \Omega} \hat{\mathrm{r}}_{dj}^{\ 2} \leq \delta$ controls the overall model complexity by keeping the weights close to zero. This constraint does not only enforce sparsity but also provides regularization on the latent representations. To the best of our knowledge, there is not matrix decomposition method that performs similar data-driven regularization. Regularizations are usually applied to individual matrix factors such as in gradient-based matrix approximation (*6*), or to the SVD singular values as in nuclear norm regularization for matrix completion (*7, 8*).

Interestingly, in the extreme case of $\alpha = 1$, our objective function is equivalent to the following optimization;

$$\min_{W\geq 0, H\geq 0} \sigma(\mathrm{W},\mathrm{H}) = \frac{1}{2} \parallel \mathrm{R} - \mathrm{WH} \parallel_F^2$$

which is, indeed, the standard Non-negative Matrix Factorization (NMF) of Lee and Seung (*9, 10*). That is, projection functions are not needed to discriminate the learning of the observed entries over the unobserved entries, as both are considered equally important. Moreover, in the extreme case of $\alpha = 0$, our objective function is equivalent to the weighted NMF used in recommendation system (*11*). In this case, the model learns only from the observed entries.

We have also explored additional variations of the method. The most obvious is applying additional L2 regularization on the low-rank matrices, as follow;

$$\min_{W\geq 0, H\geq 0} \ell(\mathrm{W},\mathrm{H}) = \frac{1}{2} \parallel \mathrm{R} - (\mathrm{WH})_R \parallel_F^2 + \frac{\alpha}{2} \parallel (\mathrm{WH})_{\neg R} \parallel_F^2 + \frac{\lambda}{2} (\parallel \mathrm{H} \parallel_F^2 + \parallel \mathrm{W} \parallel_F^2)$$

where $\lambda \geq 0$ is the regularization parameter. We also derived a multiplicative update rule for this case,

$$W \leftarrow W \circ \frac{RH^T}{((\mathrm{WH})_R + \alpha(\mathrm{WH})_{\neg R})H^T + \lambda W}$$

$$H \leftarrow H \circ \frac{W^T R}{W^T((\mathrm{WH})_R + \alpha(\mathrm{WH})_{\neg R}) + \lambda H}$$

However, we found that additional regularization on the matrices tend to interfere with the data-driven regularization. In short, $\lambda$ can hurt significantly the RMSE, even for $\alpha = 0$. This can be explained by considering that the constrain $\sum_{(d,j)\notin\Omega}(w_d {h_j}^T)^2 \leq \delta_r$ together with the L2 constrains $\sum_{(d,k)} {w_{d,k}}^2 \leq \delta_w$ and $\sum_{(k,j)} {h_{k,j}}^2 \leq \delta_h$ are conflictive constraints for different $\delta_r, \delta_w, \delta_h$ as the L2 constraints cannot take into account the unobserved entries.

We have also considered modelling biases in the data, i.e., there are some drugs that tend to produce more frequent side effects than the average, or some side effects that tend to be less frequent than the average (in this case, the average rating value over the observed entries). However, we found that although this approach works well for the Netflix problem (*7*) with well-defined distribution and fixed mean (based on the observed entries), in our problem, it is difficult to model biases due to the ill-defined distribution of the data considering both the bias from clinical trials and the uncertainty of the zeros.

#### Supplementary Note 4: Model assumption in the context of recommendation systems

*The notation here follows the notation presented in the Methods section of the main paper.*

In principle, we could have used two different models to separate the problems of side effect frequency prediction and binary prediction problems.  In fact, in the recommendation system literature, rating prediction and ranking are treated separately (*12*). In the rating prediction problem, it is assume that *all the unobserved values are missing* and the prediction must assign a rating from 1 to 5 to every missing user-movie pair. This is the standard Netflix problem that motivated the well-known Netflix competition in 2006 (*13*). In the ranking problem, the goal is to recall movies for each user in the top-N recommendations (*14, 15*). In this case, implicit representation of the data is normally used[1]. In the latter case, the use of the zero entries in the matrix for the decomposition have shown to improve the ranking performance (*14*).

Let's analyse these two models separately in the context of side effect prediction:

---

[1] In recommender system literature, a rating matrix that contains user preferences is called *explicit feedback* datasets. An example of these are the Netflix and Movielens datasets that contains rating values from 1 to 5. In this case, users provided explicit input of their preference whereas in *implicit feedback* datasets, they can be additional side information that helps to predict user preference, e.g. movie plots.

- **Ranking of drug side effects**. In recent work, we have shown that a simple matrix decomposition, when used for ranking, provides state of the art performance in binary side effect prediction problem (*16*). Given a *binary association matrix* $Y$ of n drugs and $m$ side effects, we can approximate it by the product of two low-rank matrices $Y \approx P_{nxk}Q_{nxk}$. To learn the matrices, we minimized the following loss using conjugate gradient descend (CGD):

$$\min_{P,Q} \ell_{binary}(\mathrm{P}, \mathrm{Q}) = \frac{1}{2} \parallel \mathrm{Y} - \mathrm{PQ} \parallel_F^2 + \frac{\lambda}{2}(\parallel \mathrm{P} \parallel_F^2 + \parallel \mathrm{Q} \parallel_F^2) \qquad (1)$$

where $\lambda$ is the regularization parameter that was set in cross-validation.

A standard evaluation measure employ for this problem is the AUROC (*17*). Importantly, in this case, the decomposition uses the zero values in the matrix, so the predicted distribution of scores would be *skewed* and *zero-inflated*.

- **Rating prediction of drug side effect frequencies like in movies (without the zeros).** Following the standard model used for movie recommendation system in rating prediction, we can formulate in similar fashion the frequency rating prediction. Given a matrix $\mathrm{R}$ that contains the frequency values, the model, i.e. $R \approx A_{nxl}B_{nxl}$, which only learns from the known associations (1 to 5) – and not the zeros - by minimizing the following loss:

$$\min_{A,B} \ell_{rating}(\mathrm{A}, \mathrm{B}) = \frac{1}{2} \parallel \mathrm{R} - (\mathrm{AB})_R \parallel_F^2 + \frac{\beta}{2}(\parallel \mathrm{A} \parallel_F^2 + \parallel \mathrm{B} \parallel_F^2) \qquad (2)$$

where $\beta$ is the regularization parameter that needs to be set in cross-validation. Notice here that $Y$ is the implicit version of $\mathrm{R}$. That is, trivially, $Y_{ij} = 1 \; if \; R_{ij} > 0$, or zero otherwise.

A standard evaluation measure used for this problem is the RMSE (*13*). Importantly, in this case, the predicted distribution of scores would imitate a normal distribution, similar to the one shown in Fig. S6 for $\alpha = 0$. Thus, the prediction would imply that most unobserved associations are expected to be *frequent,* because no knowledge on the zeros was introduce during the learning. This is in fact, an incorrect assumption for our problem.

Neither model in eq. (1) can effectively address the rating prediction nor model in eq. (2) can address the ranking of drug side effects. Our model represents a hybrid model between these two well-studied models in the recommender system literature.

In detail, our model de-biases the observed associations by integrating the partially observed matrix with information about the zeros. And the importance of the zeros in our model is accounted using $\alpha$. The effect of this parameter can be better understood by looking at the performance of the method in cross-validation (Fig. S8-9). We found that there is a trade-off between the AUROC and the RMSE that can be controlled with $\alpha$ (for a fixed $k$). Small confidence on the zeros minimizes the RMSE but decreases the AUROC and vice versa, high confidence on the zeros increases both, RMSE and AUROC. This can be interpreted as $\alpha$ adjusting for the unknown distribution of the drug side effects. For instance, for the extreme under-confidence case of $\alpha = 0$, the implicit hypothesis is that all the unobserved associations are missing values and that they should be drawn from the known clinical trials distribution (rating values 1 to 5): this is equivalent to the Netflix problem. In this case, although RMSE can be very small, AUROC is close to random. This is expected as none of the missing values were predicted as zeros. Moreover, for the overconfidence case of $\alpha = 1$, the implicit hypothesis is that most of the unobserved associations are indeed null values. In this case, AUROC is optimal at the expenses of an increase in the RMSE.

#### Supplementary Note 5: Matrix decomposition algorithm and convergence analysis

*The notation here follows the notation presented in the Methods section of the main paper.*

Our objective function penalizes on the zeros in the drug side effect matrix as follow;

$$\min_{W\geq 0, H\geq 0} \mathcal{L}(\mathrm{W},\mathrm{H}) = \frac{1}{2} \parallel (\mathrm{R})_R - (\mathrm{WH})_R \parallel_F^2 + \frac{\alpha}{2} \parallel (\mathrm{R})_{\neg R} - (\mathrm{WH})_{\neg R} \parallel_F^2$$

Notice that since $(\mathrm{R})_R = R$ and $(\mathrm{R})_{\neg R} = 0$, we can re-write the objective as follow:

$$\min_{W\geq 0, H\geq 0} \mathcal{L}(\mathrm{W},\mathrm{H}) = \frac{1}{2} \parallel R - (\mathrm{WH})_R \parallel_F^2 + \frac{\alpha}{2} \parallel (\mathrm{WH})_{\neg R} \parallel_F^2$$

We shall now prove the following theorem following the procedure in (*18*).

**Theorem 3.1**. *The functional $\mathcal{L}(W,H)$ converges to a local minimum under the update rule*

$$W \leftarrow W \circ \frac{RH^T}{((\mathrm{WH})_R + \alpha(\mathrm{WH})_{\neg R})H^T}$$

$$H \leftarrow H \circ \frac{W^T R}{W^T((\mathrm{WH})_R + \alpha(\mathrm{WH})_{\neg R})}$$

*Proof.* From the theory of constrained optimization, we know that to prove Theorem 3.1, we need to show that at convergence, the solution satisfies the well-known Karush-Khun-Tucker (KKT) complementary condition (*18*):

$$\left(\frac{\partial \mathcal{L}}{\partial \mathrm{W}}\right)_{dk} W_{dk} = 0, \left(\frac{\partial \mathcal{L}}{\partial \mathrm{H}}\right)_{kj} H_{kj} = 0 \qquad (1)$$

The gradients can be written in matrix form;

$$\frac{\partial \mathcal{L}}{\partial \mathrm{W}} = -\,(\mathrm{R} - (\mathrm{WH})_R)\, H^T + \alpha(\mathrm{WH})_{\neg R} H^T \qquad (2)$$

$$\frac{\partial \mathcal{L}}{\partial \mathrm{H}} = -\,W^T(R - (\mathrm{WH})_R) + \alpha W^T (\mathrm{WH})_{\neg R} \qquad (3)$$

At local minimum, $\mathrm{W} = W^*$ and $\mathrm{H} = H^*$ must satisfy the KKT conditions in (1). Replacing the gradients (2) and (3) in (1);

$$\left(\mathrm{R}\, H^T\right)_{dk} W_{dk} - \left( ((\mathrm{WH})_R + \alpha(\mathrm{WH})_{\neg R})H^T\right)_{dk} W_{dk} = 0 \qquad (4)$$

$$\left(W^T R\right)_{kj} H_{jk} - \left( W^T ((\mathrm{WH})_R + \alpha (\mathrm{WH})_{\neg R})\right)_{dk} H_{kj} = 0 \quad (5)$$

Also, at convergence, the multiplicative rule is,

$$W^*_{dk} = W^*_{dk} \frac{\left(RH^{*T}\right)_{dk}}{\left((W^* H^*)_R + \alpha (W^* H^*)_{\neg R})\, H^{*T}\right)_{dk}}$$

$$H^*_{dk} = H^*_{dk} \frac{\left(W^{*T} R\right)_{dk}}{\left(W^{*T} (W^* H^*)_R + \alpha (W^* H^*)_{\neg R})\right)_{dk}}$$

which are identical to our KKT conditions in (4) and (5). Therefore, the algorithm converges to a local minimum. ■

### Supplementary Figures

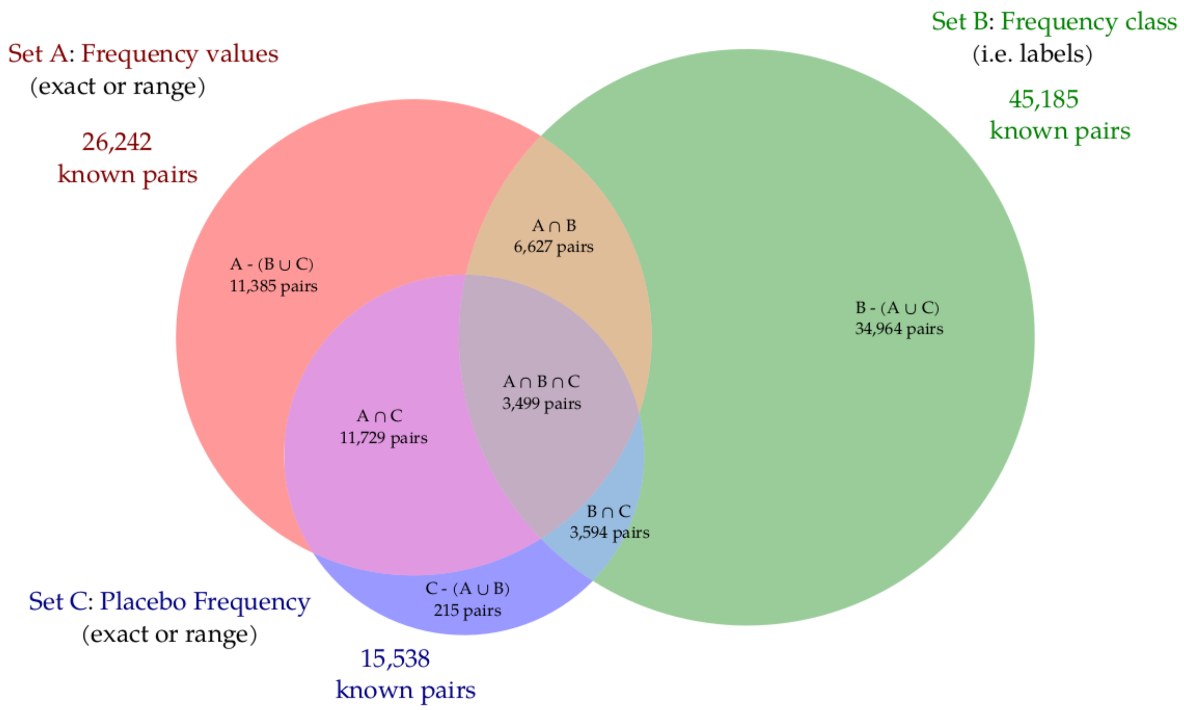


**Figure S1.** Venn diagram depicting the different formats for the drug side effect frequencies in SIDER 4.1. In total, 68,514 pairs were found with frequency data. The frequency dataset was divided into three overlapping sets. set A: contains drug exact (i.e. 1%) and range frequency (i.e. 2-5%); set B: contains frequency class (i.e. very rare); and set C contains the exact and range placebo frequency.  The size of the circles is proportional to the number of drug-side effect pairs in each set.

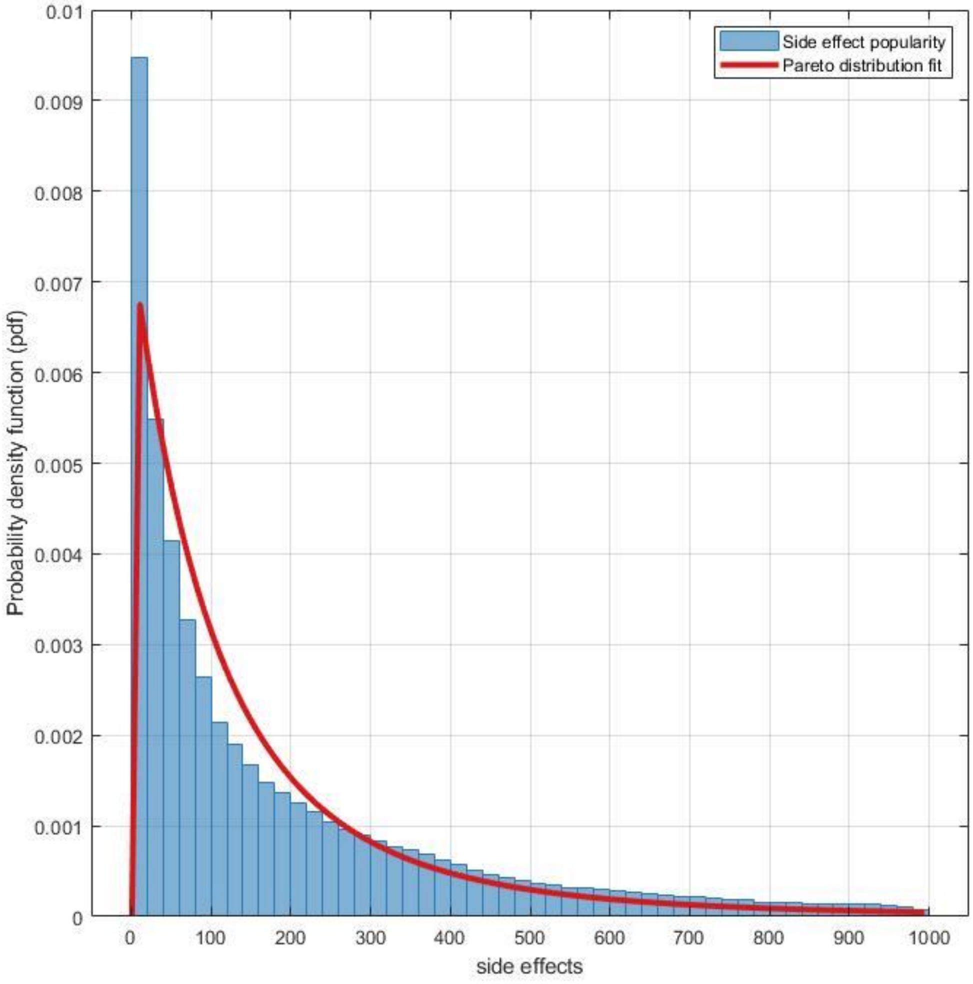


**Figure S2. Distribution of side effect popularity and tail data modelling using Generalized Pareto distribution**. Side effects are ordered in decreasing order of popularity. This includes 759 drugs and 994 side effects.  We used the matlab built-in function gpfit to estimate the tail index parameter (k = 0.2960) and the scale parameter (sigma = 135.996).

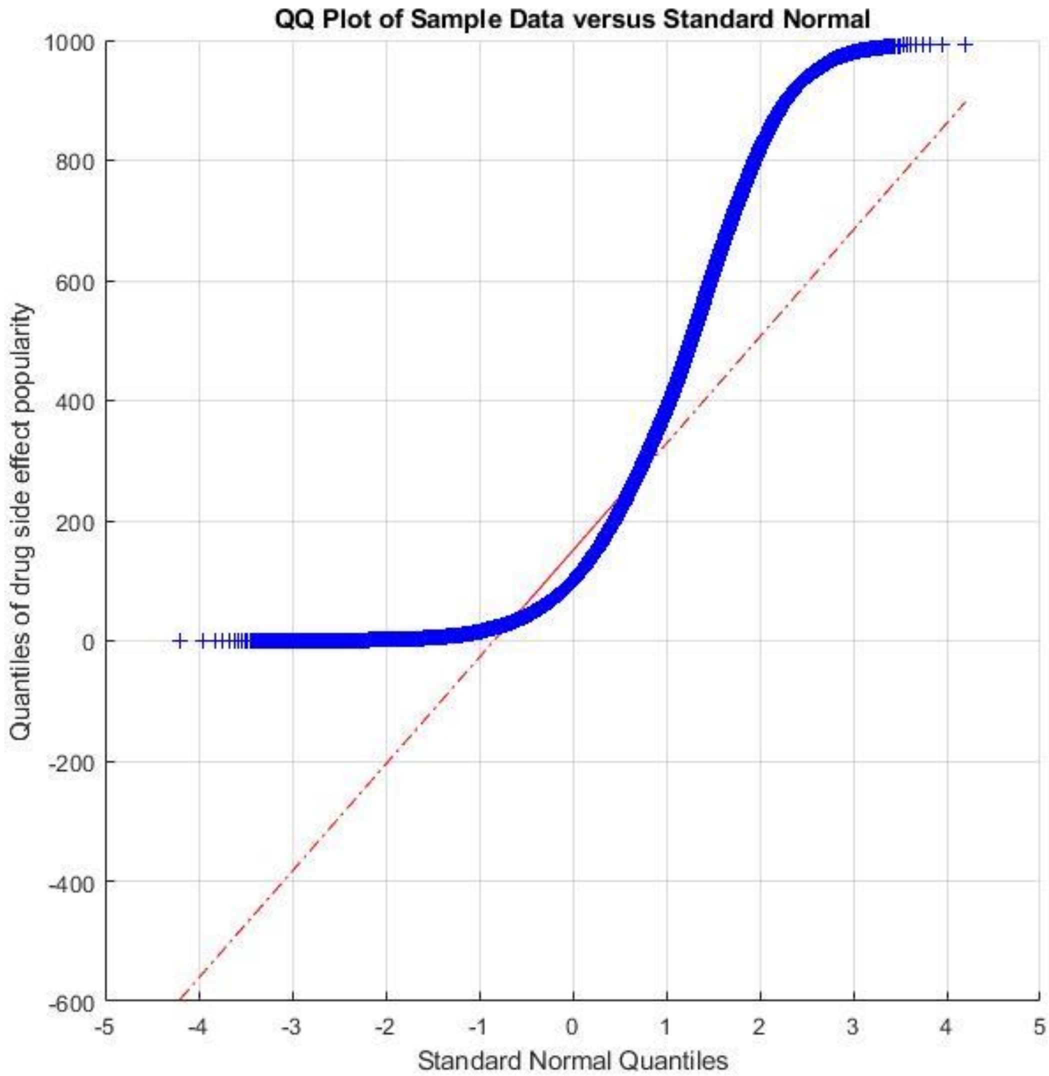


**Figure S3. QQ plot of side effects popularity distribution.** This includes 759 drugs and 994 side effects. The plot shows the quantile-quantile distribution of the data versus the theoretical quantile value from a normal distribution. The kurtosis is a measure of how outlier-prone the distribution is. The kurtosis of the drug side effects is 5.04, in comparison the kurtosis of a normal distribution is 3.

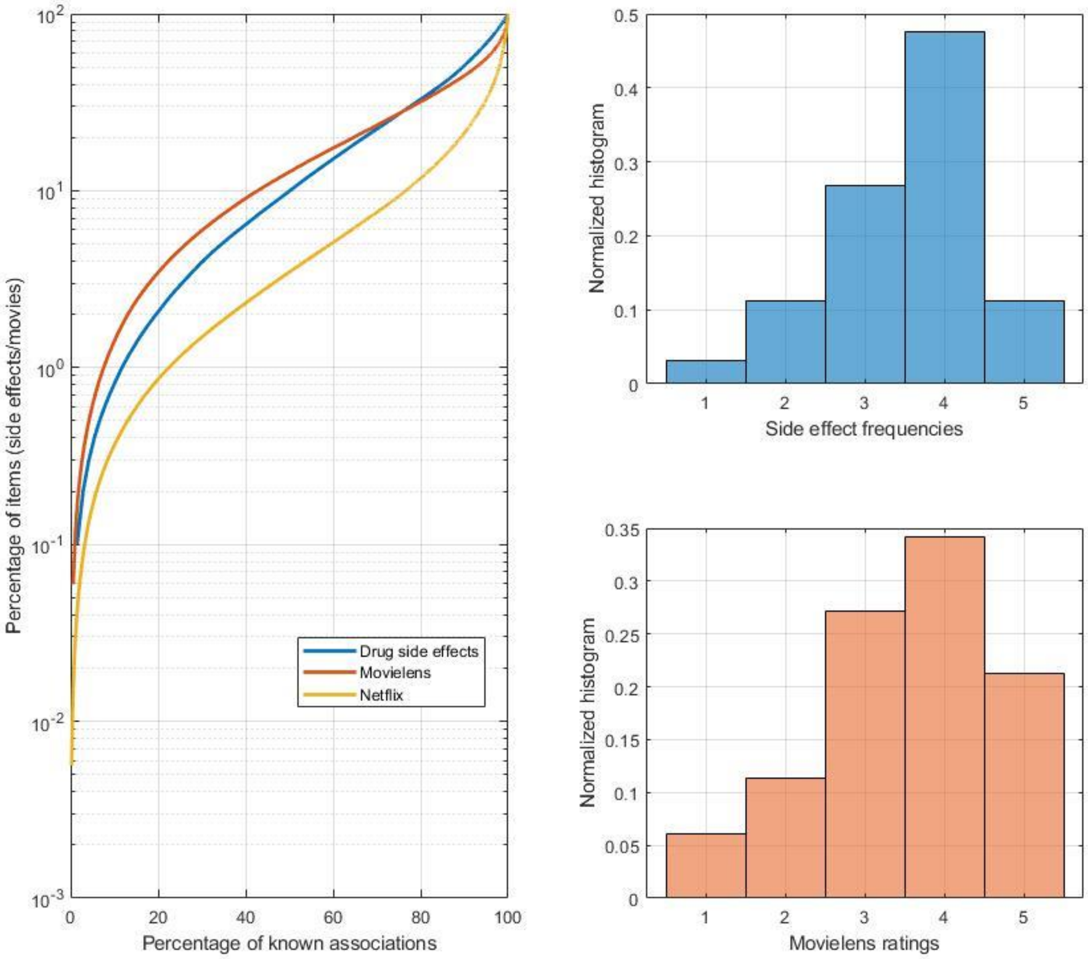


**Figure S4. The heavy-tailed distribution of drug side effects and of two popular movie datasets.** *(Left)* Side effects and movies (items) are ordered according to popularity, most popular at the bottom. Side effects and movie datasets tend to have few popular items containing more than 20% of the associations (usually known as *short-head*). However, most items reside in the *long-tail* of the distribution, populated with items with fewer known associations. The Movielens dataset contains 943 users and 1682 movies with 100K associations (~6.3% density). The Netflix dataset contains 480,189 users and 17,770 movies with 100M associations (~1.17% density). The density of the Movielens dataset is more comparable to our dataset of drug side effects (~4.96% density). *(Right)* Distribution of rating values for drug side effect frequency and the rating values in the Movielens dataset. The distribution of frequency values comes from a normal distribution (Chi-square goodness-of-fit Significance, $p < 2.2251e-308$) and it is very similar to the distribution of ratings in Movielens (Kolmogorov-Smirnov Significance, $p < 2.51e-233$).

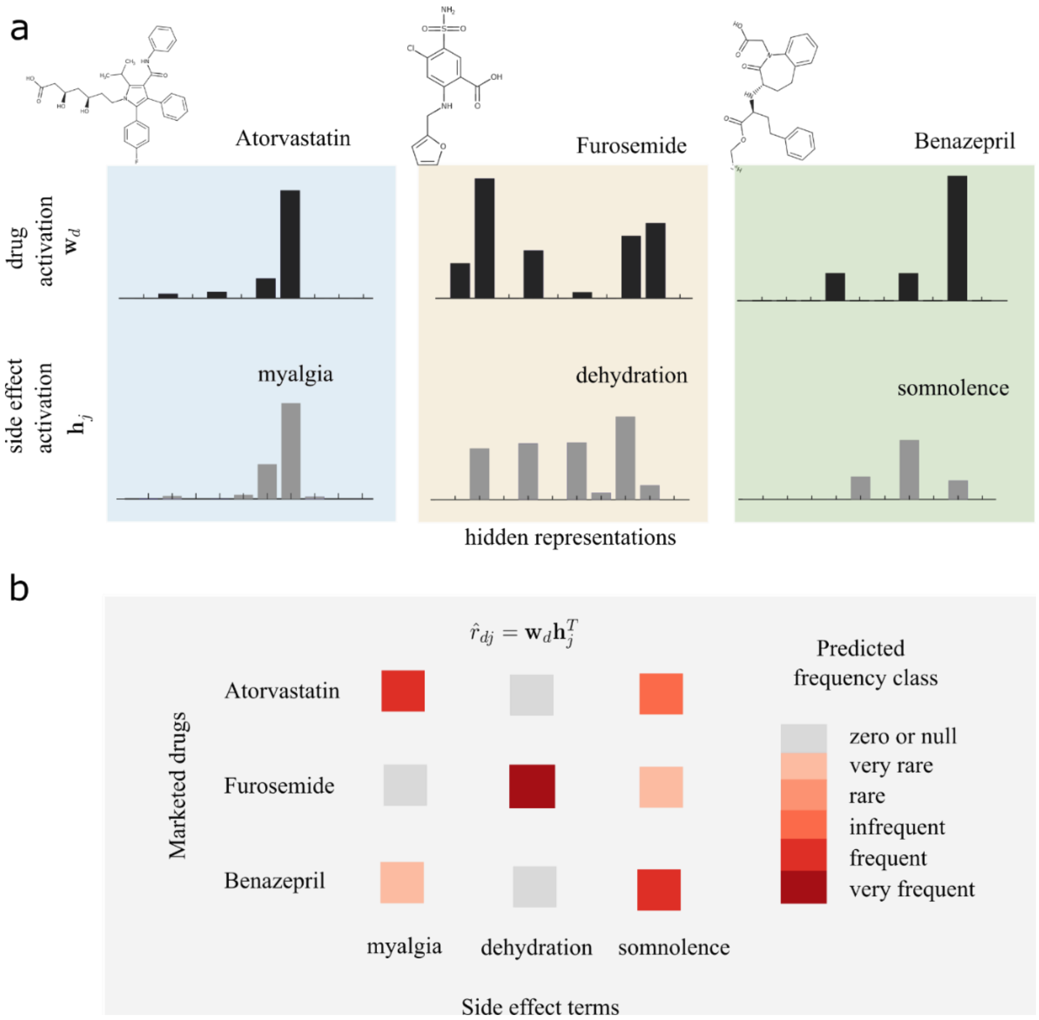


**Figure S5. Drug side effect frequency prediction from drugs and side effects signatures**. In this simulated example, **(a)** three different drugs and three different side effect activation patterns (signatures) are shown. Let's assume that the cholesterol-lowering agent (atorvastatin) causes muscle pain (myalgia), the diuretic agent (furosemide) causes dehydration and the hypertensive drug (benazepril) causes somnolence. We should expect that their activation patterns over a set of latent representations (feature vector) is then very similar. Thus, the most correlated drug-side effect patterns are shown in similar background color **(b)** To generate a specific drug side effect frequency estimation, the dot product between both activation patterns is computed. In this illustration, the square's colors indicate the predicted frequency class.

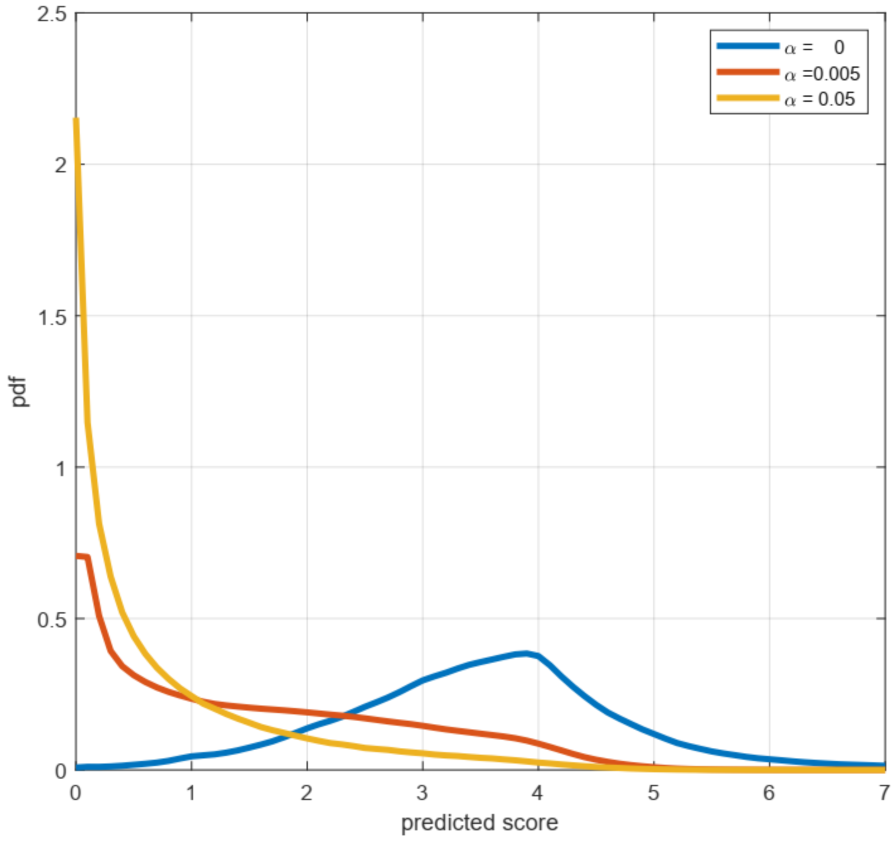


**Figure S6. Distribution of predicted scores in our model for different values of $\alpha$.** The figure shows the probability density estimation of the predicted scores in our model for three different values of $\alpha = (0, 0.005, 0.05)$ and a value of $k = 10$ for all the cases. The PDF shows that when $\alpha = 0$, the model only learns from the known frequency classes, i.e. 1 to 5, and therefore, it incorrectly assumes that the zeros are not a possible outcome. In the other cases ($\alpha \neq 0$), the model learns from the zeros with small confidence. The distribution becomes long-tailed and zero-inflated, and the priors on the classes change accordingly. Hence, rarer side effects, associated to lower scores, are now more likely than the more frequent ones, associated to higher scores. This has the important effect of de-biasing the observed distribution of clinical trials.

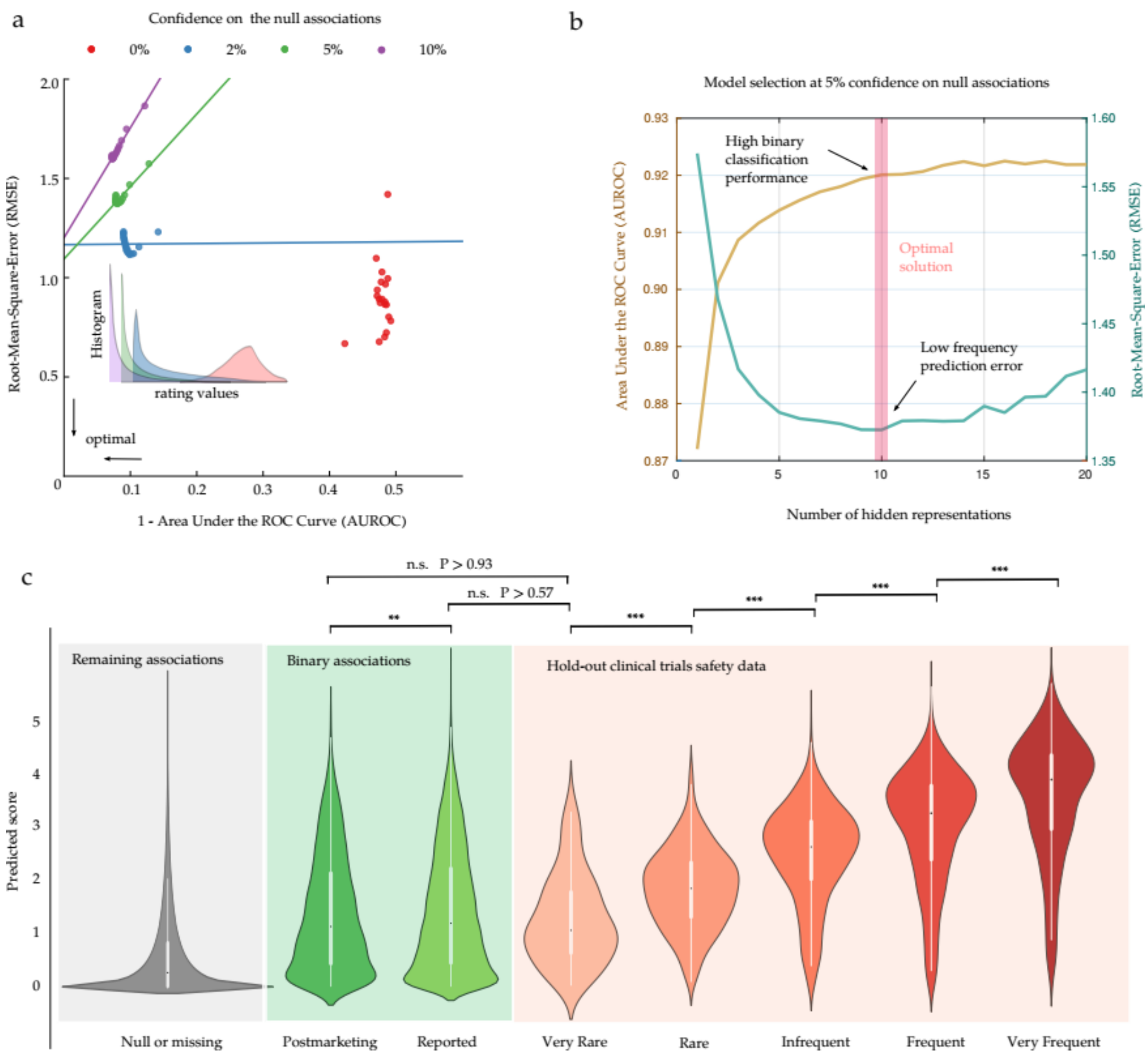


**Figure S7. Model selection and prediction of drug side effects frequency. (a)** Training performance of RMSE versus 1-AUROC for varying values of $\alpha = (0, 0.02, 0.05, 0.1)$ (shown as percentage in red, blue, green and purple) for different model complexity (each point for a $k \in [1, \ldots, 20]$). $\alpha = 5\%$ gives close to optimal AUROC with good RMSE. *Inset*. Predicted distribution of drug side effects frequency for all the pairs for each of the four models at a fixed $k = 10$. **(b)** Selection of the optimal number of hidden representations based on the RMSE-AUROC trade-off. **(c)** Predicted score for the binary associations: post market and reported associations (green) and a hold-out test set with known frequency classes from clinical trials (red, true classes are indicated in the x-axis). The predicted score for the remaining associations is also shown (grey). Our method can accurately discriminate between all the different frequency classes: rare vs very rare (Significance, $p < 2.80x10^{-12}$), infrequent vs rare (Significance, $p < 1.31x10^{-40}$), frequent vs infrequent (Significance, $p < 3.45x10^{-51}$) and very frequent vs frequent (Significance, $p < 9.00x10^{-26}$). Interestingly, not significant differences were found between the prospective sets and the scores for the very rare side effects (Significance, $p > 0.936$ for post marketing and $p > 0.578$ for reported side effects), but significant differences were found when the prospective set was chosen at random (Significance, $p < 1.08x10^{-25}$). Significance levels between the scores are indicated with asterisks ($p \leq 0.001$, ***), ($p \leq 0.01$, **), and n.s. stands for non-significant.

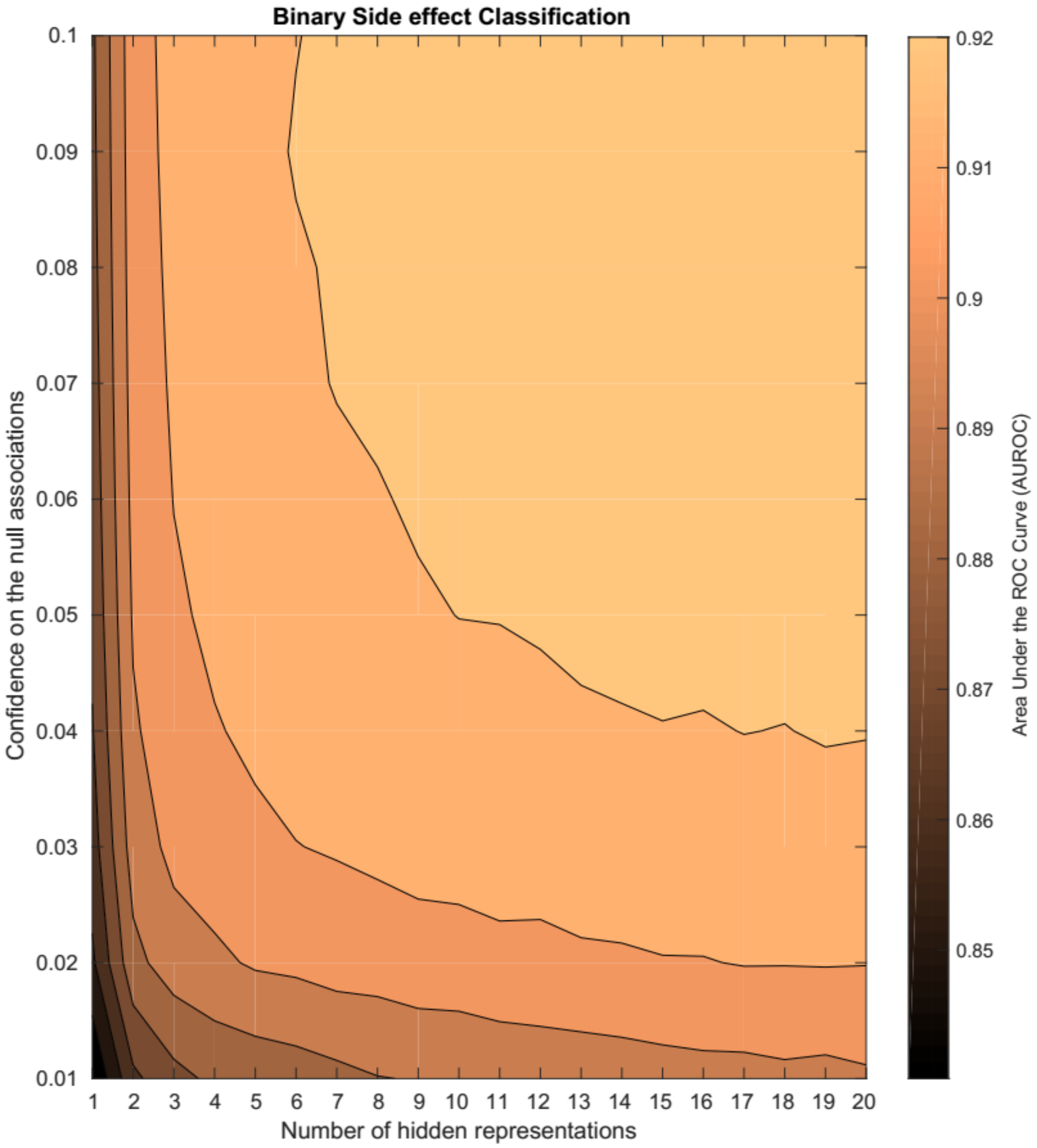


**Figure S8. Contour plot of mean AUROC of the 10FCV training performance for the binary side effect classification problem (true vs false).** The higher the AUROC, the better we can identify true associations from unknown ones. The performance is divided for clarity in nine contour levels for varying values of the number of hidden representations ($k$) and the confidence on the zeros ($\alpha$).

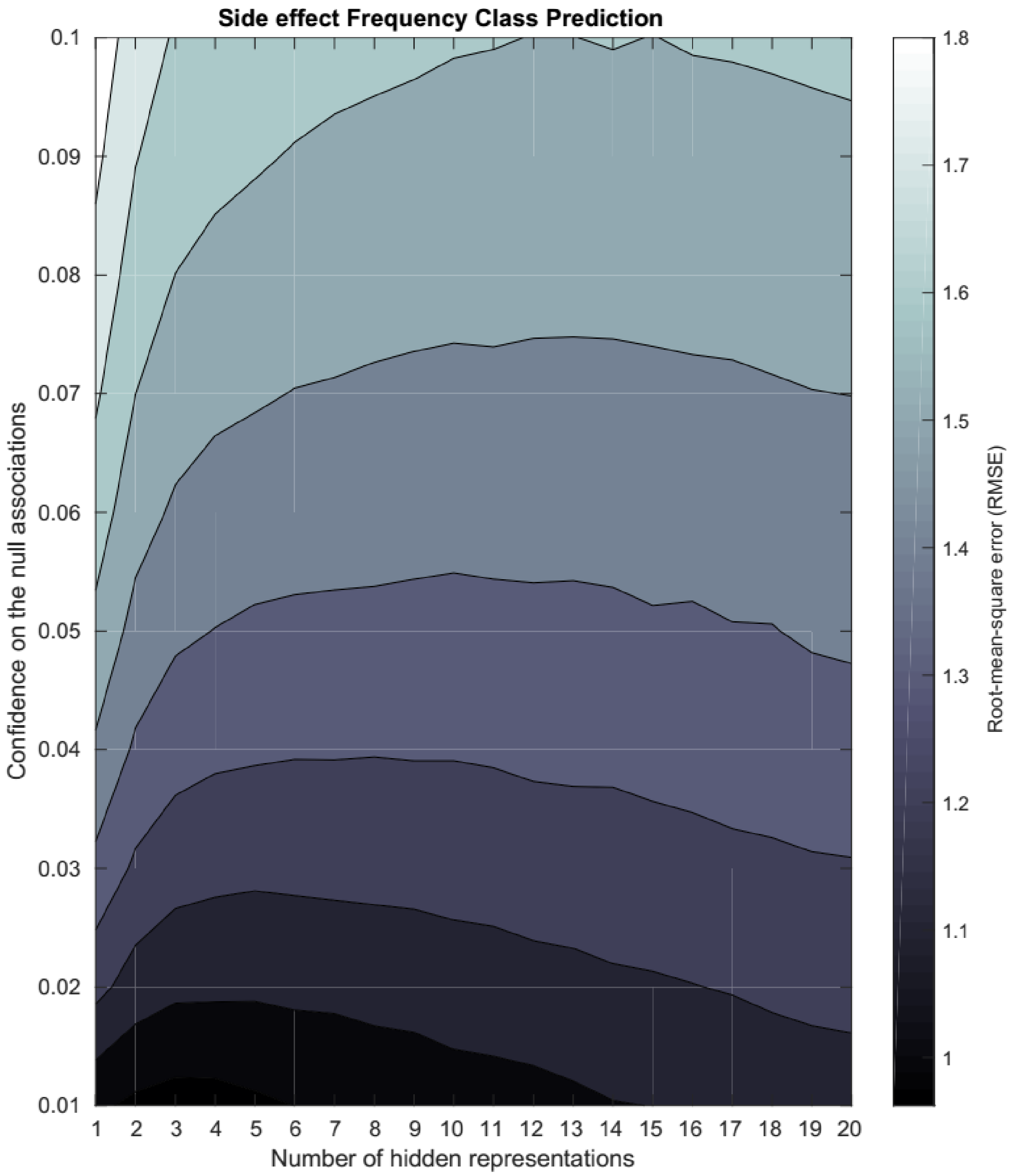


**Figure S9. Contour plot of mean RMSE of the 10FCV training performance for the side effect frequency rating prediction problem (1, 2, 3, 4, 5).** The smaller the RMSE, the better we can predict the true frequency value of the drug side effects. The performance is divided for clarity in nine contour levels for varying values of the number of hidden representations ($k$) and the confidence on the zeros ($\alpha$).

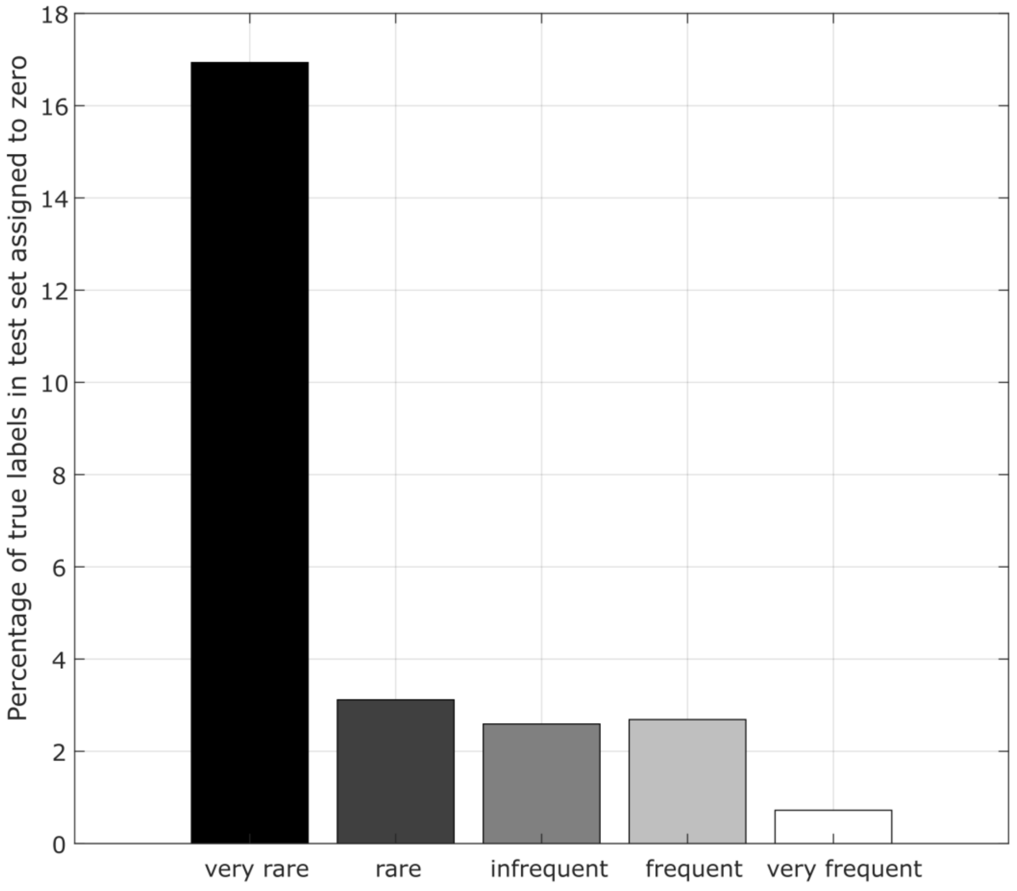


**Figure S10. Percentage of true labels in the hold-out test set assigned to zero for each of the side effect frequency classes**. Around 15% of the very rare drug side effects were assigned to zeros, likely driven by the few numbers of labels in the dataset. However, less than 3% of the associations belonging to the other classes were incorrectly classified as false drug side effect.

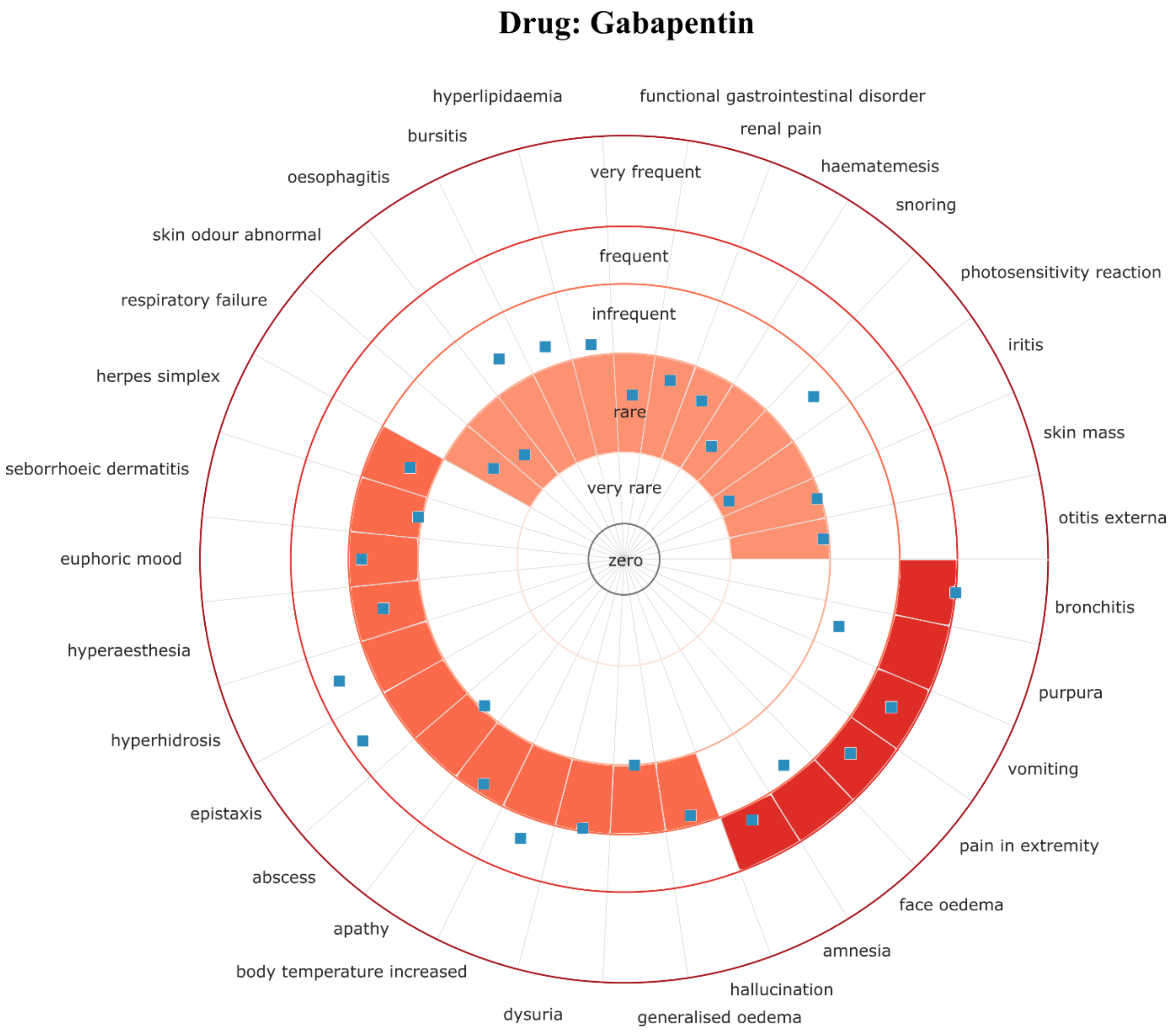


**Fig. S11. Predictions for all the side effects produced by the drug Gabapentin from the held-out test set.** The thirty-one predictions in the test set for the anticonvulsant drug Gabapentin are shown around polar plots, each in a dedicated sector. Concentric circles between frequency classes correspond to thresholds learned by maximum likelihood. The correct class for each association is coloured in each circular sector while predicted scores are shown as blue squares.

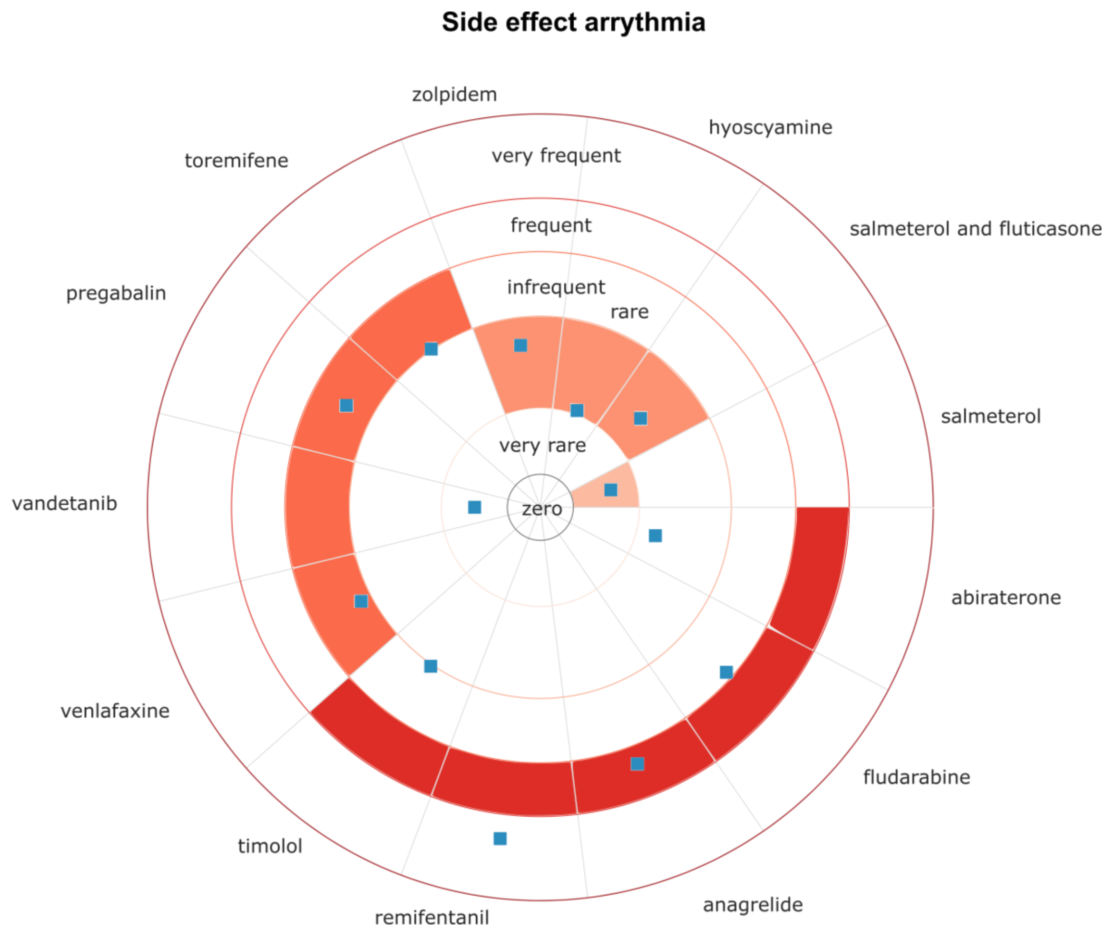


**Fig. S12. Predictions for all the drugs that produce arrhythmia from the held-out test set.** The thirteen predictions in the test set for the cardiovascular side effect arrhythmia are shown around polar plots, each in a dedicated sector. Concentric circles between frequency classes correspond to thresholds learned by maximum likelihood. The correct class for each association is colored in each circular sector while predicted scores are shown as blue squares.

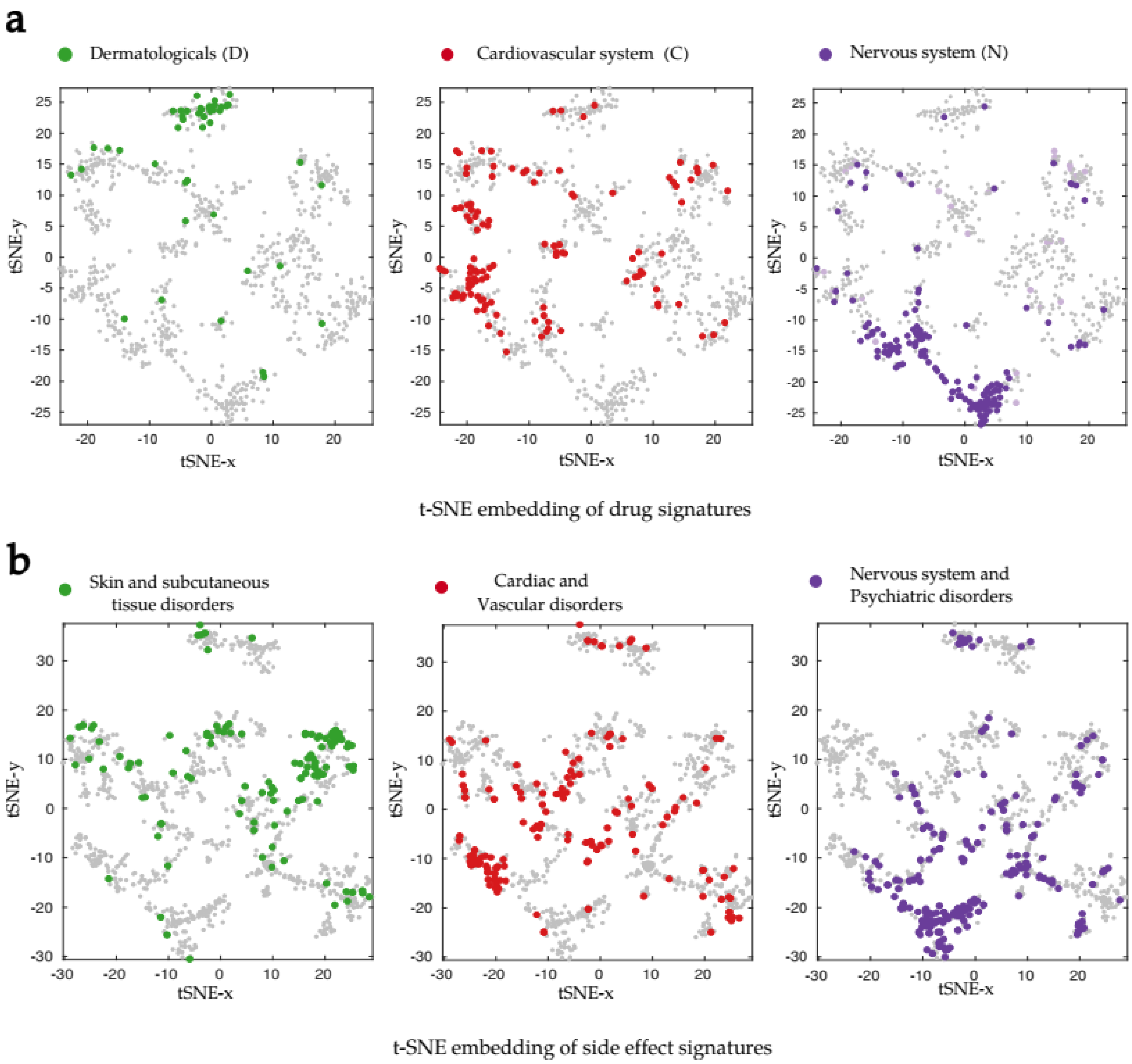


**Figure S13. Embedding of drug and side effect signatures in 2D space using t-SNE. (a)** In the drug signature embedding, each point represents a drug colored according to their main Anatomical, Therapeutic and Chemical (ATC) category (*left*) Dermatologicals; (*mid*) Cardiovascular system (C) and; (*right*) Nervous system (N) drugs. Remaining drugs are shown in grey in each plot. **(b)** In the side effect signature embedding, each point represents a side effect colored according to their main Medical Dictionary for Regulatory Activities (MedDRA) classification of disorders. (*left*) Sin and subcutaneous tissue disorders, (*mid*) Cardiac and Vascular disorders and (*right*) Nervous system and Psychiatric Disorders. Remaining side effects are shown in grey in each plot.

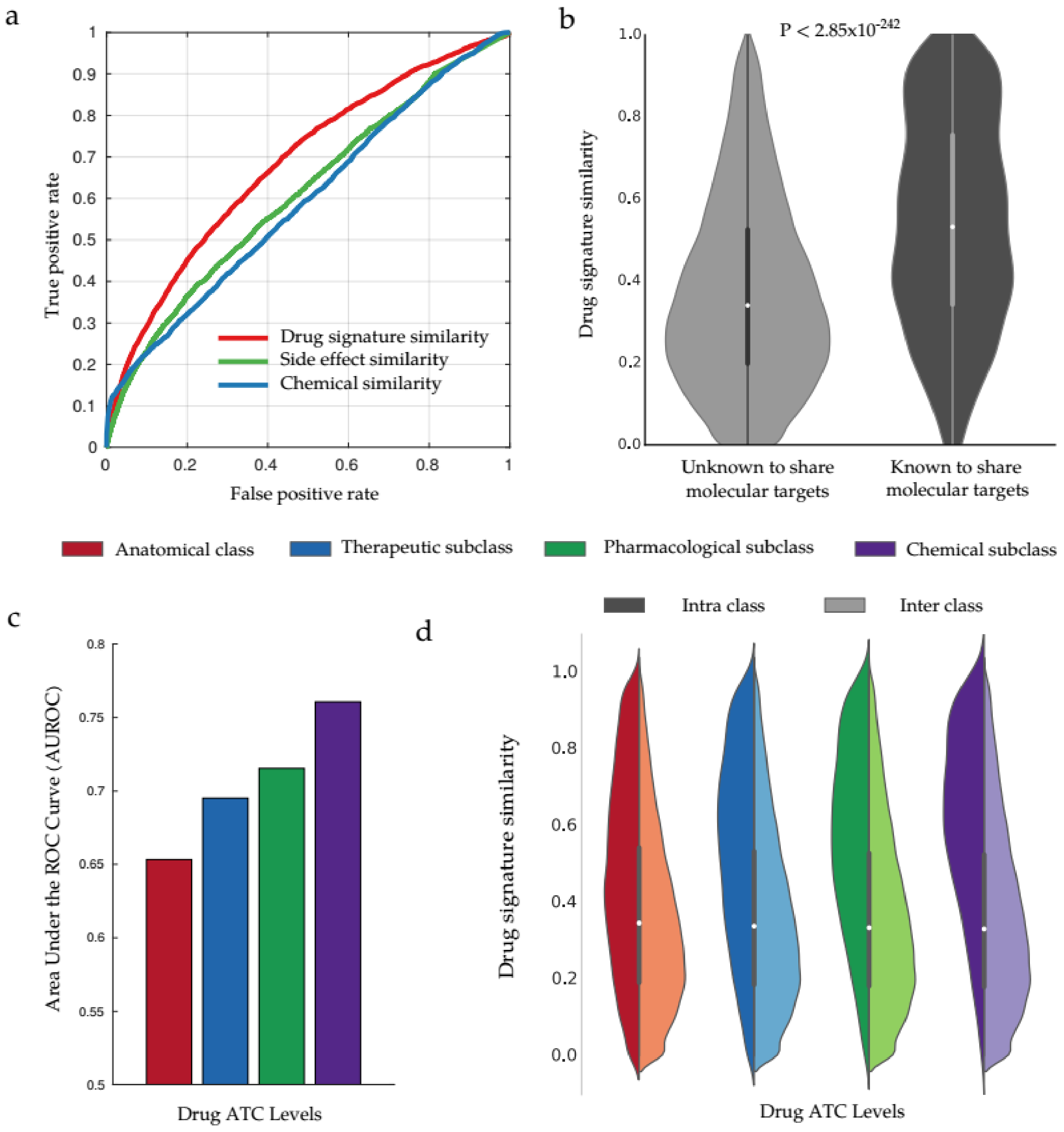


**Figure S14. Predicting shared protein targets and shared clinical activity using drug signature similarities. (a)** ROC curve representing the ability of the drug signature similarity, side effect similarity and 2D Tanimoto chemical similarity scores to predict which pairs of drugs share targets. **(b)** Distribution of the drug signature similarity for the pairs of drugs known and not known to shared protein targets. **(c)**

Area Under the Receiver Operating Characteristic Curve (AUROC) representing the ability of the drug signature similarity to predict whether which pairs of drugs share Anatomical, Therapeutic and Chemical (ATC) category for each of the different levels of drug activity in the taxonomy: anatomical (red), therapeutic (blue), pharmacological (green) and chemical (purple). **(d)** Comparative distribution of drug signature similarity for the intraclass (dark color), *i.e.* pairs that share ATC category, versus the interclass (light color), i.e. pairs not known to share ATC in the indicated level.

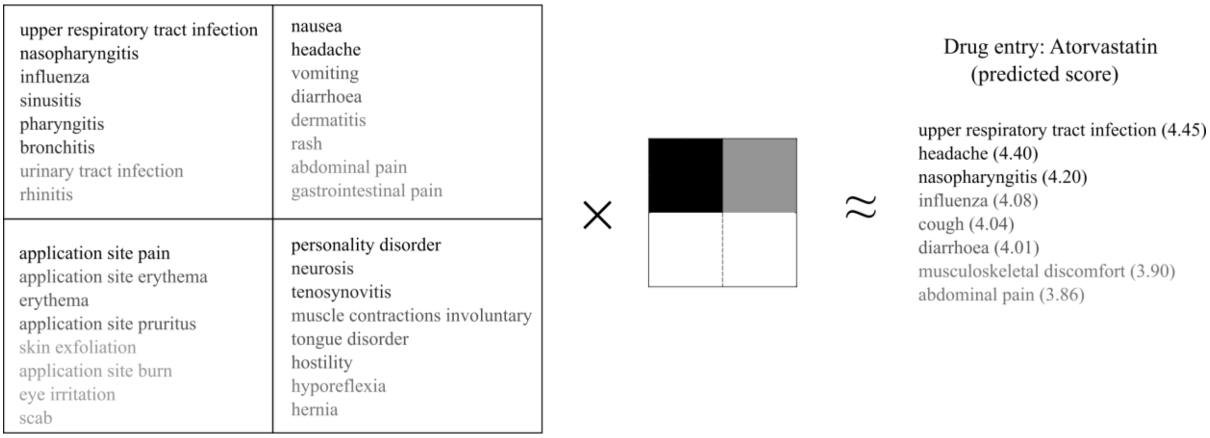


**Figure S15. Anatomical/physiological relationships between side effects.** For the drug side effect frequency matrix R we used the decomposition algorithm to obtain the two matrices W and H. Upper left, four of the 10 latent representations obtained (rows of H). Side effect terms for each latent representation was obtained according to their weight to the specific latent variable. The darkness in the side effect term indicates the strength of the connection to the latent variable. Right, the eight most frequently predicted side effects for the cholesterol-lowering agent Atorvastatin (predicted scores shown in parenthesis). This frequency was approximated by a superposition that gave weight to the upper two latent representations, and none to the lower two, as shown by the four shaded squares in the middle indicating the activations in W. It can be seen that the model learn the anatomical relationships between side effects, as they tend to be group by disorder-category, e.g. respiratory system related side effects are strongly connected to the darkest latent variable in the middle.

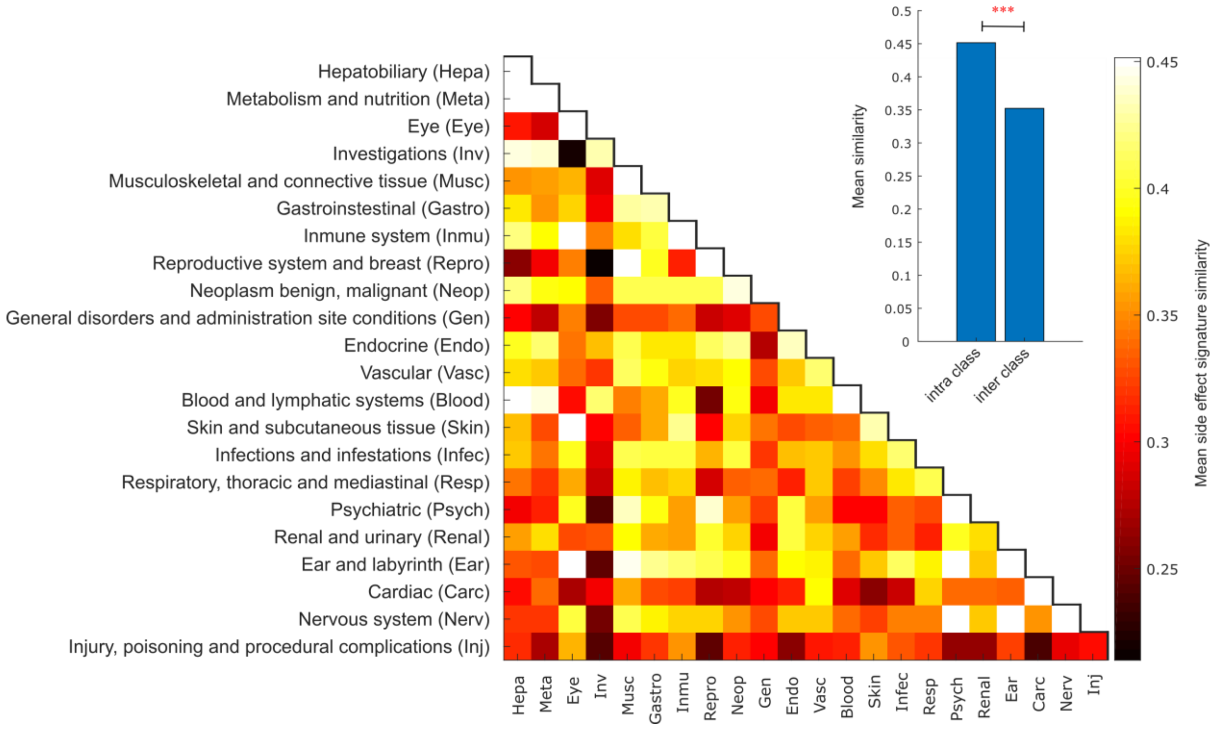


**Fig. S16. Side effect signatures encodes side effect phenotypes.** Each (x, y) tile represents, for each main Medical Dictionary for Regulatory Activities (MedDRA) classification of disorders, the mean similarity of side effect pairs where one side effect belong to category x and the other to category y. The value ranges from 0.21 (Reproductive systems - Investigations) to 0.58 (Psychiatric – Psychiatric). The colours range between the minimum mean similarity and 0.45, with all values above 0.45 (In the diagonal: 0.49 (Hepa), 0.55 (Eye), 0.57 (Repro), 0.49 (Blood), 0.58 (Psych), 0.54 (Carc), 0.47 (Nerv)) set to 0.45. Inset: the average intra-class similarity is significantly higher than the average inter-class similarity (t-test p-value < 4.37e-16).

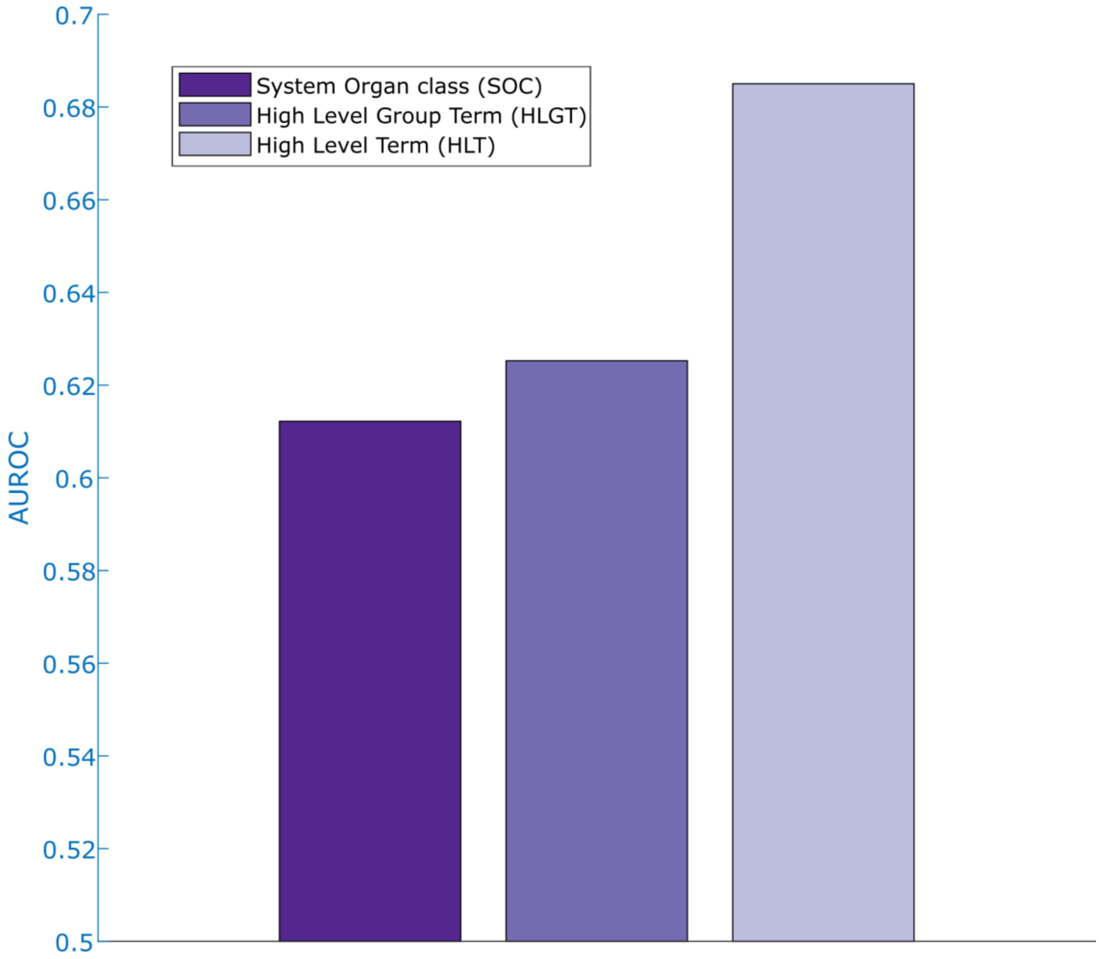


**Figure S17. Predicting share side effect anatomical/physiological categories for different levels of the MedDRA taxonomy using side effect signatures similarity**. Level 1 or System Organ class (SOC): 57,076 side effects that share and 436,445 do not. Level 2 or High Level Group Term (HLGT): 12,097 share and 481,424 do not. Level 3 or High Level Term (HLT): 2,312 share and 491,209 do not.

### Supplementary Tables

**Table S1.** Mapping of frequency values to frequency classes and to rating values.

| Frequency class | Intervention cohort-clinical trials frequency | Assigned rating value |
|---|---|---|
| very frequent | More than 10% | 5 |
| frequent | 1 to 10% | 4 |
| infrequent | 0.1 to 1% | 3 |
| rare | 0.01 to 0.1% | 2 |
| very rare | Less than 0.01% | 1 |
| zeros* | 0% | 0 |
| *Zeros are not usually provided in safety datasets as only true positive cases are reported. | | |

**Table S2.** Number of drugs for each main drug Anatomical, Therapeutic and Chemical (ATC) Category. There are 14 main categories that classifies drugs by their clinical use. Drug categories are ordered by the number of drugs in the gold-standard. 676 drugs (89.06%) belongs to one category only, whereas the remaining can belong to more than one category. The number of known associations with side effect frequency is also shown together with the average premarketing side effect frequency.

| **Top ATC drug category** | **Number of drugs** | **Number of known associations w/ side effect frequency** | **Average frequency rating value** |
|---|---|---|---|
| Nervous system (N) | 142 | 15,470 | 3.183 |
| Antineoplastic and immunomodulating agents (L) | 129 | 7,286 | 4.042 |
| Cardiovascular system (C) | 111 | 2,993 | 3.666 |
| Antiinfectives for systemic use (J) | 105 | 4,135 | 3.620 |
| Alimentary tract and metabolism (A) | 83 | 2,108 | 3.660 |
| Genito urinary system and sex hormones (G) | 50 | 1,632 | 3.706 |
| Sensory organs (S) | 47 | 1,491 | 3.471 |
| Respiratory system (R) | 46 | 987 | 3.811 |
| Dermatologicals (D) | 45 | 1,604 | 3.671 |
| Various (V) | 31 | 609 | 3.568 |
| Blood and blood forming organs (B) | 31 | 839 | 3.639 |
| Musculo squeletal system (M) | 30 | 1,275 | 3.519 |
| Systemic hormonal preparations, insulins (H) | 18 | 465 | 3.948 |
| Antiparasitic products, insecticides and repellents (P) | 10 | 235 | 3.906 |

**Table S3.** Number of side effect terms for each main Side effect MedDRA category of disorders. The categories are ordered on decreasing number of side effects. Side effect terms could belong to more than one main category. Most side effects belong to a one or two category of disorders (by 59.65%).

| Top MedDRA category of disorders | Number of side effects | Number of known associations w/ side effect frequency | Average side effect frequency |
|---|---|---|---|
| Nervous system disorders | 136 | 5,868 | 3.527 |
| Skin and subcutaneous tissue disorders | 119 | 4,167 | 3.378 |
| Gastrointestinal disorders | 116 | 5,913 | 3.715 |
| Vascular disorders | 97 | 3,725 | 3.342 |
| Respiratory, thoracic and mediastinal disorders | 96 | 3,604 | 3.590 |
| General disorders and administration site conditions | 92 | 4,463 | 3.807 |
| Psychiatric disorders | 88 | 3,409 | 3.499 |
| Infections and infestations | 84 | 2,964 | 3.611 |
| Eye disorders | 67 | 1,336 | 3.206 |
| Reproductive system and breast disorders | 61 | 1,390 | 3.370 |
| Metabolism and nutrition disorders | 60 | 2,127 | 3.524 |
| Cardiac disorders | 58 | 2,806 | 3.469 |
| Injury, poisoning and procedural complications | 58 | 870 | 3.410 |
| Investigations | 57 | 1,794 | 3.670 |
| Musculoskeletal and connective tissue disorders | 57 | 2,585 | 3.570 |
| Renal and urinary disorders | 48 | 1,544 | 3.240 |

| | | | |
|---|---|---|---|
| Blood and lymphatic system disorders | 43 | 1,466 | 3.283 |
| Immune system disorders | 42 | 1,249 | 2.611 |
| Neoplasms benign, malignant and unspecified (incl cysts and polyps) | 25 | 206 | 3.053 |
| Hepatobiliary disorders | 24 | 483 | 2.687 |
| Endocrine disorders | 24 | 563 | 3.115 |
| Ear and labyrinth disorders | 13 | 506 | 3.328 |
| Social circumstances | 3 | 70 | 3.571 |
| Surgical and medical procedures | 2 | 59 | 3.271 |
| Pregnancy, puerperium and perinatal conditions | 2 | 13 | 2.384 |
| Congenital, familial and genetic disorders | 1 | 5 | 3.400 |
